# GID E3 ligase supramolecular chelate assembly configures multipronged ubiquitin targeting of an oligomeric metabolic enzyme

**DOI:** 10.1101/2021.04.07.436316

**Authors:** Dawafuti Sherpa, Jakub Chrustowicz, Shuai Qiao, Christine R. Langlois, Laura A. Hehl, Karthik Varma Gottemukkala, Fynn M. Hansen, Ozge Karayel, Susanne von Gronau, J. Rajan Prabu, Matthias Mann, Arno F. Alpi, Brenda A. Schulman

**Affiliations:** Department of Molecular Machines and Signaling, Max Planck Institute of Biochemistry, Martinsried, 82152, Germany; Department of Proteomics and Signal Transduction, Max Planck Institute of Biochemistry, Martinsried, 82152, Germany

## Abstract

To achieve precise cellular regulation, E3 ubiquitin ligases must be configured to match substrate quaternary structures. Here, by studying the yeast GID complex, mutation of which is ***<u>G</u>***lucose-***<u>I</u>***nduced ***<u>D</u>***egradation deficient, we discover supramolecular chelate assembly as an E3 ligase strategy for targeting an oligomeric substrate. Cryo EM structures show that to bind the tetrameric substrate fructose-1,6-bisphosphatase (Fbp1), two otherwise functional GID E3s assemble into a 20-protein Chelator-GID^SR4^, which resembles an organometallic supramolecular chelate. The Chelator-GID^SR4^ assembly avidly binds multiple Fbp1 degrons and positions Fbp1 so that its protomers are simultaneously ubiquitylated at lysines near its allosteric and substrate binding sites. Significantly, key structural and biochemical features -including capacity for supramolecular assembly - are preserved in the human ortholog, the CTLH E3. Based on our integrative structural, biochemical and cell biological data, we propose that higher-order E3 ligase assembly generally underlies multipronged targeting, capable of simultaneously incapacitating multiple protomers and functionalities of oligomeric substrates.

## INTRODUCTION

Cells respond to changes in nutrient availability by rapidly adapting their metabolic pathways (Tu and McKnight, 2006; Zaman et al., 2008; Zhu and Thompson, 2019). Shifts in metabolic enzyme activities are typically achieved through simultaneous regulation at seemingly every conceivable level. First, metabolite-responsive transcriptional programs activate pathways to maximally use available nutrients, and to repress those rendered unnecessary or counterproductive. Second, the catalytic activities of metabolic enzymes are subject to tight allosteric control. Metabolic enzymes are often oligomers, with inter-subunit interactions cooperatively regulating catalysis by transmitting signals from metabolite binding or protein modifications (Koshland, 1963a, b; Monod et al., 1963). Third, in eukaryotes, undesired metabolic activities are often terminated by ubiquitin-mediated proteolysis (Nakatsukasa et al., 2015).

Degradation is typically controlled by recognition of proteins as substrates for ubiquitylation by E3 ligases. However, little is known about whether, and if so how, E3 ligases are specifically tailored for oligomeric assemblies of metabolic enzymes. One of the first identified targets of a eukaryotic nutrient-dependent suppression pathway, budding yeast fructose-1,6-bisphosphatase (Fbp1), is an oligomer (Chiang and Schekman, 1991). Fbp1 is a gluconeogenic enzyme essential for yeast growth on non-fermentable carbon sources. As shown 40 years ago, following a shift from carbon starved to rich conditions, gluconeogenesis becomes unnecessary and energetically costly, and accordingly, Fbp1 activity and expression are both immediately curtailed (Gancedo, 1971; Schork et al., 1994a; Schork et al., 1994b; Schork et al., 1994c; Schork et al., 1995). Notably, Fbp1’s enzymatic activity depends on its homotetrameric assembly; inter-subunit interactions allosterically respond to metabolites serving as substrates or allosteric inhibitors (Ke et al., 1990a; Ke et al., 1990b). Metabolic conditions also regulate Fbp1 levels. The switch to glycolytic conditions induces ubiquitin-mediated degradation of Fbp1 by the multiprotein E3 ligase aptly named “GID”, for ***<u>G</u>***lucose-***<u>I</u>***nduced ***<u>D</u>***egradation deficient (Braun et al., 2011; Chiang and Schekman, 1991; Menssen et al., 2012; Regelmann et al., 2003; Santt et al., 2008; Schork et al., 1994a; Schork et al., 1995). Although the GID E3 is conserved across eukaryotes and regulates important physiology, its biochemical mechanisms and targets are best characterized in budding yeast.

Much like well-studied multiprotein E3 ligases such as Anaphase-Promoting Complex/Cyclosome or Cullin-RING ligases, GID is not a singular complex (Barford, 2020; Liu and Pfirrmann, 2019; Melnykov et al., 2019; Qiao et al., 2020; Rusnac and Zheng, 2020; Watson et al., 2019). The constituents of various GID assemblies, and how they achieve regulation is beginning to emerge. Under carbon starvation conditions, a core but inactive multiprotein complex is assembled, termed GID^Ant^ due to its production in anticipation of a future shift in conditions eliciting GID E3 activity (Qiao et al., 2020). The GID^Ant^ assembly comprises a scaffold of subunits Gid1, Gid5 and Gid8, and a catalytic module comprising the largely homologous proteins Gid2 and Gid9 wherein Gid2 displays a hallmark E3 ligase RING domain. However, GID^Ant^ lacks substrate-binding subunits. GID^Ant^ is converted into an active E3 ligase upon binding to one of two mutually exclusive substrate receptors induced in response to different environmental perturbations (Qiao et al., 2020). A switch from gluconeogenic to glycolytic conditions triggers production of the substrate receptor subunit Gid4, which associates with GID^Ant^ to form a GID^SR4^ complex that contains all the fundamental elements of a functional E3 ligase: substrate binding capability and RING-catalyzed ubiquitin transferase activity (Menssen et al., 2018; Qiao et al., 2020; Santt et al., 2008). GID^SR4^ is an Pro/N-degron pathway E3 ligase, whereby Gid4 recruits an N-terminal proline in peptide-like degron motifs in the substrates Fbp1, malate dehydrogenase (Mdh2) and isocitrate lyase (Icl1) (Chen et al., 2017; Dong et al., 2018; Hämmerle et al., 1998). Ubiquitin transfer is catalyzed by Gid2 and Gid9, which together activate the cognate E2∼ubiquitin intermediate with the E2, Ubc8 (aka Gid3; “∼” here refers to reactive thioester bond linking the E2 catalytic cysteine and the ubiquitin C-terminus) (Schüle et al., 2000). The mechanisms revealed by cryo EM structures of GID^Ant^ and GID^SR4^ were validated by mutational effects on ubiquitylation of Mdh2 *in vitro* and glucose-induced Fbp1 degradation *in vivo* (Qiao et al., 2020). However, Fbp1 ubiquitylation has not been reconstituted *in vitro* using defined GID E3 ligase components.

Several observations pointed toward a distinct, but elusive GID E3 “complex” mediating Fbp1 regulation. Perplexingly, Fbp1 degradation *in vivo* depends on another protein, Gid7 (Regelmann et al., 2003). Gid7 was identified based on its association with other Gid proteins (Menssen et al., 2012; Santt et al., 2008). However, Gid7 is dispensable for GID^SR4^ assembly, and for glucose-regulated stability of another GID E3 ligase substrate (Negoro et al., 2020). Moreover, Gid7 is evolutionarily conserved across eukaryotes (Francis et al., 2013; Kobayashi et al., 2007). Mammals even have two orthologs, WDR26 and MKLN1, which both co-immunoprecipitate subunits of the so-called “CTLH E3” multiprotein complex that contains the mammalian orthologs of yeast GID^SR4^ subunits (Boldt et al., 2016; Francis et al., 2013; Kobayashi et al., 2007; Lampert et al., 2018; Liu and Pfirrmann, 2019; Salemi et al., 2017). The CTLH E3, named based on the preponderance of CTLH domains (in Gid1, Gid2, Gid7, Gid8 and Gid9 and their orthologs across evolution), has intrinsic E3 ligase activity (Lampert et al., 2018; Liu and Pfirrmann, 2019; Maitland et al., 2019). Although Pro/N-degron substrates of the CTLH E3 are yet to be identified, human Gid4 structurally superimposes with yeast Gid4 assembled into GID^SR4^ and likewise preferentially binds peptides with an N-terminal proline (Dong et al., 2020; Dong et al., 2018; Qiao et al., 2020). Physiological importance of the CTLH E3 is underscored by developmental defects in flies, fish, frogs, and mammals arising from genetic deletions or mutations of its subunits (Han et al., 2016; Javan et al., 2018; Nguyen et al., 2017; Pfirrmann et al., 2015; Soni et al., 2006; Wei et al., 2019; Zavortink et al., 2020; Zhen et al., 2020). Yet, how Gid7 or its human orthologs regulate GID and CTLH E3 ligase complexes is unknown.

Here, through identifying Gid7 orthologs as GID/CTLH E3 assembly factors, we discover the transformation of a multiprotein E3 ligase complex (GID^SR4^) into a supramolecular “Chelator”-GID^SR4^ complex. Studying yeast proteins, we show that Gid7 nucleates a 20-protein supramolecular chelate E3 ligase assembly that is specifically tailored for Fbp1’s quaternary structure. Our data reveal supramolecular chelate assembly of a pre-existing functionally competent E3 ligase complex as a structural and functional principle to achieve multipronged ubiquitin targeting tailored to the structure and function of an oligomeric substrate.

## RESULTS

### Fbp1 ubiquitylation involves a supramolecular GID E3 assembly

We considered that there might be a distinct substrate-specific GID E3 assembly beyond the structurally-determined GID^SR4^, because the Gid7 protein, not visualized previously, is required for timely glucose-induced regulation of Fbp1, but not of the GID^SR4^ substrate Mdh2 *in vivo* (Negoro et al., 2020; Regelmann et al., 2003; Santt et al., 2008). Thus, to examine effects on degradation, we employed the promoter reference technique. This allows monitoring degradation of an exogenously expressed protein (here C-terminally Flag-tagged Fbp1) while normalizing for effects on transcription and translation (Chen et al., 2017; Oh et al., 2017). Notably, Gid7 is essential for Fbp1 degradation in this assay, with effects of its deletion comparable to that of removing Gid4, the subunit responsible for binding Fbp1’s degron sequence (Figure 1A).

**Figure 1.**
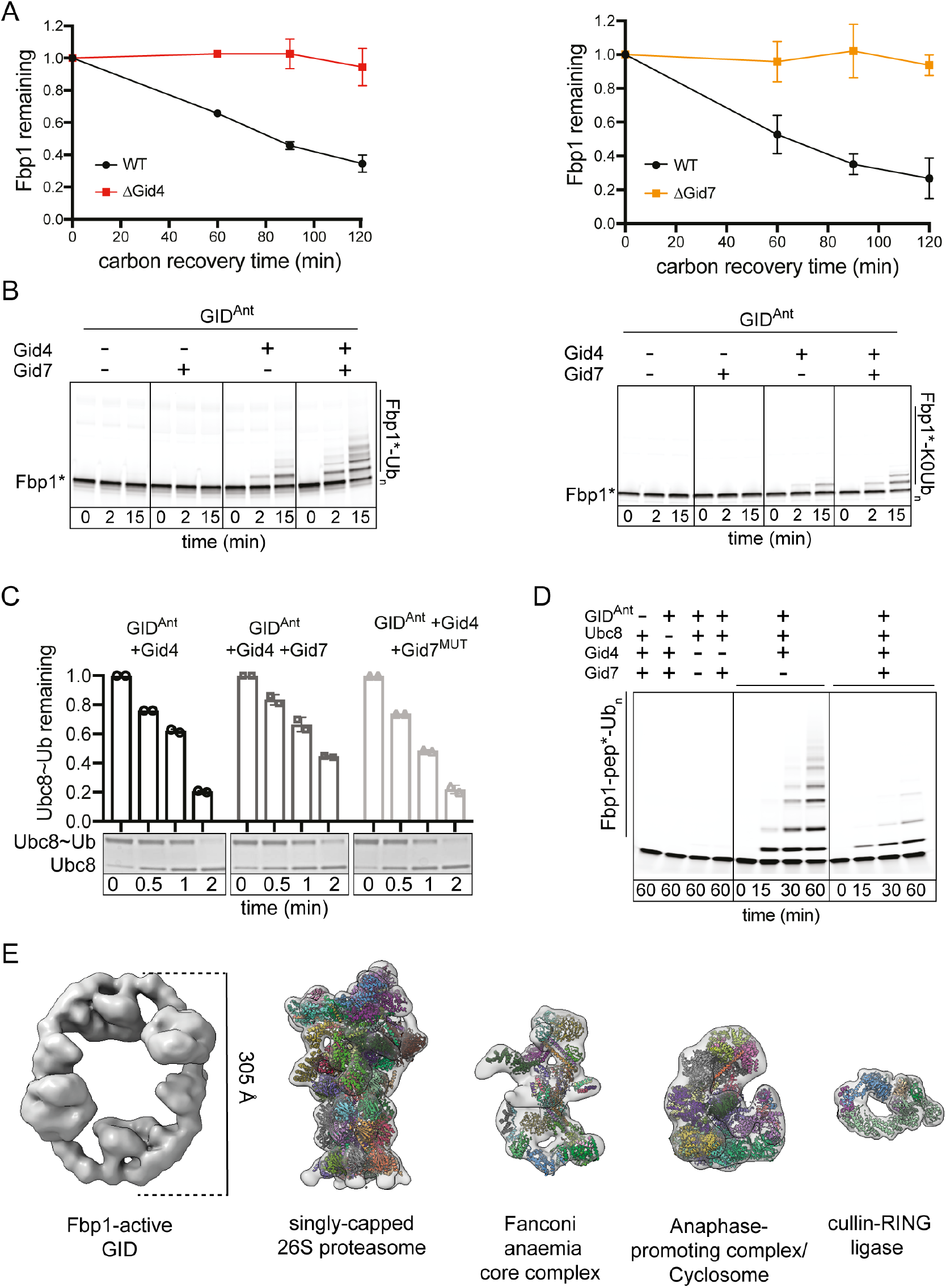
Distinct GID E3 ligase assembly specified by Gid7 (See also Figure S1) A. Assays testing roles of Gid7 (right) and substrate receptor Gid4 (left) on glucose-induced degradation of Fbp1 *in vivo*, quantified using the promoter reference technique. Relative protein levels in WT, Gid4 and Gid7 KO yeast strains, after terminating ethanol starvation and initiating glucose recovery (timepoint 0), were calculated as the ratio of exogenously expressed Fbp1-3xFLAG compared to the reference protein DHFR-3xHA expressed from the same promoter. Error bars represent SD (n>3). B. Assays testing roles of Gid7 and Gid4 on Fbp1 ubiquitylation *in vitro*. Reactions were performed with WT ubiquitin (left) or its lysineless variant (K0, right). GID^Ant^ is the complex lacking Gid4 and Gid7. GID^SR4^ is the complex formed by adding Gid4 substrate receptor to GID^Ant^. Progress of the reaction was followed by visualizing C-terminally fluorescently-labelled Fbp1 (denoted as Fbp1*). C. Effect of Gid7 on substrate-independent GID ubiquitin transferase activity monitored in a pulse-chase format. During the pulse, the thioester-linked Ubc8∼ubiquitin intermediate was generated, and the reaction was quenched. In a chase, discharge of ubiquitin from Ubc8 was initiated by adding free lysine and GID^SR4^ with or without Gid7 (Gid7^WT^) or its mutant lacking LisH-CTLH-CRA motif, Δ1-285 (Gid7^MUT^). Progress of the reaction was visualized by Coomassie-stained non-reducing SDS-PAGE. The percentage of remaining non-discharged Ubc8∼Ub was normalized against timepoint 0 and plotted. Error bars represent SD (n=2). D. Assays testing roles of Gid7 on ubiquitylation of monomeric, model peptide substrate *in vitro*. The peptide substrate (Fbp1-pep) comprises the N-terminal Fbp1 degron and an acceptor lysine connected via a flexible linker, and a C-terminal fluorescein. Assay was performed using GID^SR4^ with and without addition of Gid7. Control lanes show the dependence of ubiquitylation activity on GID^SR4^, E2 Ubc8 and substrate receptor Gid4. E. Low resolution cryo EM structure of GID complex active toward Fbp1 (GID^SR4^ + Gid7) compared to other multi-protein complexes. Models fit into low-pass filtered maps shown for comparison are singly-capped 26S proteasome (PDB: 5MPB, EMDB: EMD-3536), Fanconi Anaemia core complex (PDB: 6SRI, EMDB: EMD-10290), Anaphase promoting complex/Cyclosome (PDB: 5L9T, EMDB: EMD-3433) and cullin-RING E3 ubiquitylation complex (PDB 6TTU, EMDB: EMD-10585).

A series of biochemical assays reconstituting GID E3 ligase activity *in vitro* revealed multimodal Gid7-dependent transformation of GID^SR4^ into distinct E3 ligase assembly specifying Fbp1 ubiquitylation. First, adding Gid7 to GID^SR4^ (generated by adding the substrate receptor Gid4 to a GID^Ant^ complex) greatly increased Fbp1 ubiquitylation (Figure 1B, left). This activation depends on the substrate receptor subunit, because GID^Ant^ was inactive towards Fbp1 even with Gid7. Comparing reactions with WT ubiquitin or a “K0” version lacking lysines that cannot form polyubiquitin chains indicated that adding Gid7 increases substrate consumption, number of Fbp1 sites modified, and number of ubiquitins in polyubiquitin chains (Figure 1B, right). Second, the activation upon adding Gid7 was specific to the Fbp1 substrate. Adding Gid7 actually suppressed the instrinsic E3 ligase activity of GID^SR4^, as monitored by effects on ubiquitin transfer from a pre-formed Ubc8∼ubiquitin intermediate to free lysine in solution (Figure 1C). Binding of Fbp1’s degron per se is insufficient to overcome this inhibition, because Gid7 likewise subdued ubiquitylation of a model peptide substrate wherein Fbp1’s degron sequence, PTLV (Chen et al., 2017), is connected to a lysine acceptor through an intervening flexible linker (Figure 1D). Thus, we hypothesized that Gid7 might be impacting the GID E3 assembly. Indeed, examining complex formation *in vitro* by size exclusion chromatography, we observed a striking shift to a high molecular weight assembly when Gid7 was added to GID^SR4^ (Figure S1A).

To gain further insight, we visualized the high-molecular weight GID E3 complex by cryo EM (Figure 1E). A reconstruction at 13 Å resolution showed a remarkable structure: an exterior oval supporting several inward pointing globular domains. Strikingly, the longest exterior dimension of 305 Å is roughly comparable to that of a singly-capped 26S Proteasome, 1.3 times that of the multiprotein Fanconi Anemia E3 ligase complex and 1.5 times that of the APC/C mediating polyubiquitylation (Brown et al., 2016; Chen et al., 2016; Haselbach et al., 2017; Lander et al., 2012; Schweitzer et al., 2016; Shakeel et al., 2019; Wehmer et al., 2017). Unlike these compact assemblies, however, this GID complex displays a behemoth hollow center -with interior edges of 270 and 130 Å in the longest and shortest dimensions, respectively – larger than an 8-protein CRL E3 complex ubiquitylating a substrate (Baek et al., 2020).

### A supramolecular Chelator-GID^SR4^ E3 assembly encapsulates the tetrameric Fbp1 substrate

The organization of the GID assembly was gleaned from comparing cryo EM maps of complexes comprising selected Gid subunits (Figure 2A). Two copies of the previously defined “GID^SR4^” E3 ligase structure were readily visualized in the larger assembly (Qiao et al., 2020). This version of GID^SR4^, used for the previously published high resolution cryo EM structure and affinity purified based on a tag at the N-terminus of Gid1 was a stoichiometric complex of Gid1-Gid2-Gid4-Gid5-Gid8-Gid9 (Qiao et al., 2020). However, the version used for biochemical assays and herein was affinity purified based on a tag at the C-terminus of Gid8. The latter version contains one additional protomer each of Gid1 and Gid8, which could be observed in a cryo EM map of this GID^SR4^ at subnanometer resolution, but structural heterogeneity of the additional density precluded its visualization upon further refinement. The extra Gid1-Gid8 subcomplex is well-defined and present in two copies in the Gid7-dependent assembly. Finally, the large oval-shaped GID assembly displayed yet additional density we interpreted as four copies of Gid7, in two dimers. Purified Gid7 formed a dimer on its own, as measured by SEC-MALS (Figure S1B). The data altogether indicated that the large GID assembly is a 1.5 MDa eicosamer composed of four Gid1 : two Gid2 : two Gid4 : two Gid5 : four Gid7 : four Gid8 : two Gid9 protomers.

**Figure 2.**
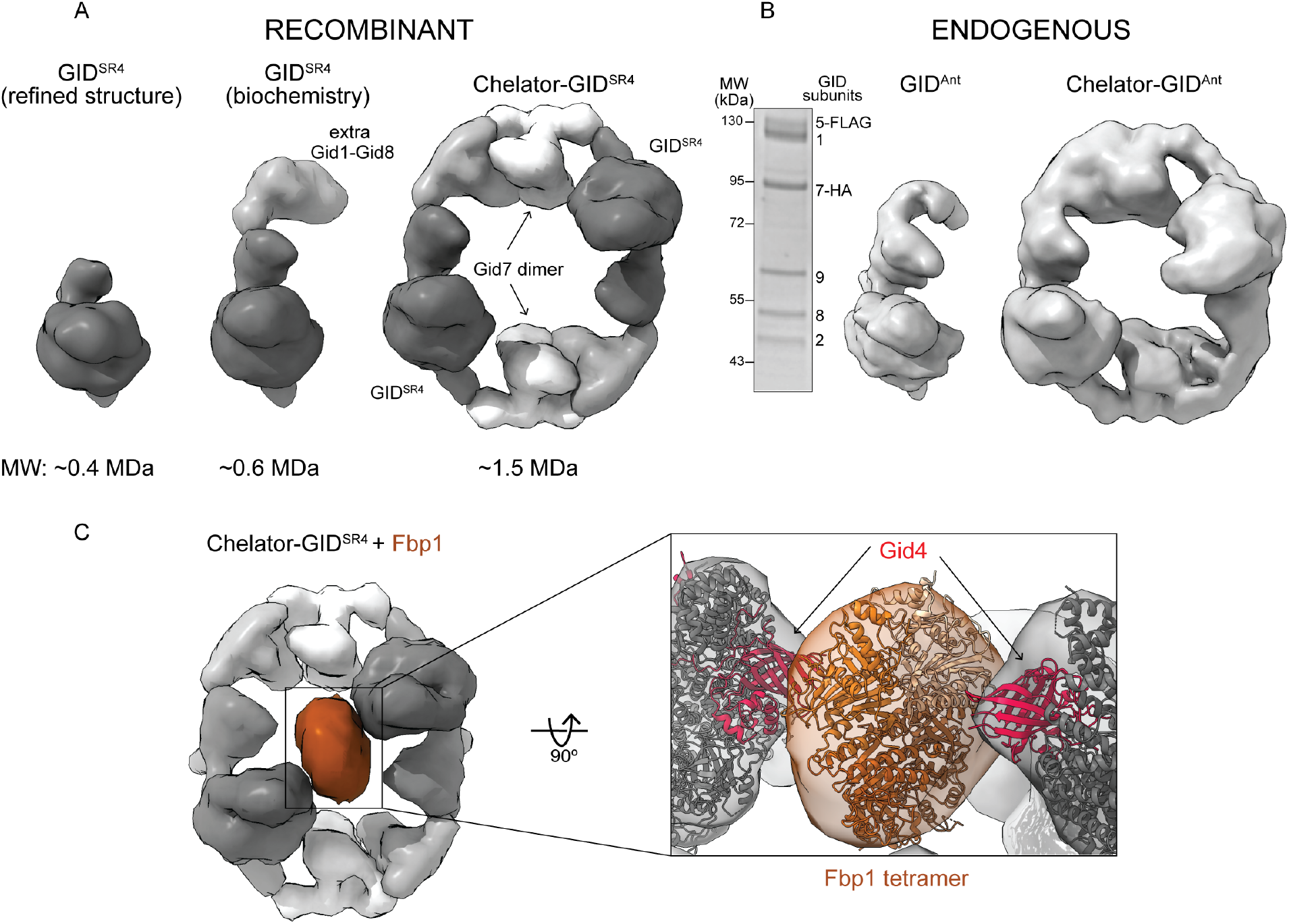
Multidentate capture of Fbp1 tetramer by Chelator-GID^SR4^ assembly (See also Figure S2) A. Cryo EM maps and molecular weights of recombinant GID assemblies. Structurally-determined GID^SR4^ (left, low pass-filtered, dark gray, EMD 10327, 6SWY.PDB) is a stoichiometric complex of Gid1, Gid8, Gid5, Gid4, Gid2 and Gid9. Purification conditions used herein include an additional Gid1-Gid8 subcomplex (gray) bound to GID^SR4^ (middle, taken for biochemical assays). Oval higher-order “Chelator-GID^SR4^” assembly includes Gid7 dimers (right, white). B. Coomassie-stained SDS-PAGE (left) and cryo EM maps of endogenous yeast GID^Ant^ (middle) and Chelator-GID^Ant^ (right) assemblies (prepared by FLAG-IP of yeast lysates with Gid5 3xFLAG-tagged and Gid7 HA-tagged at their endogenous loci). C. Cryo EM map of Chelator-GID^SR4^ (gray) bound to Fbp1 tetramer (brown). Close-up shows 2 red Gid4 protomers (modeled from 6SWY.PDB) simultaneously contacting docked Fbp1 crystal structure.

We sought to determine if this GID assembly might be formed *in vivo*. Prior studies did (Santt et al., 2008) or did not (Qiao et al., 2020) observe Gid7 cosedimenting with other GID proteins in density gradients. This raised the possibility that, like the equally giant 26S proteasome, some subunits or regulatory partners may be prone to dissociation, for example based on lysis conditions (Leggett et al., 2002). Cryo EM data of endogenous GID purified from yeast grown under carbon starvation using mild lysis conditions yielded 3D reconstructions corresponding to the recombinant assemblies with and without Gid7, at 14.2 and 9.5 Å resolution, respectively (Figures 2B and S1C).

Why is the ubiquitin ligase for Fbp1 so giant and hollow? We hypothesized that such an assembly would form in order to accomodate the substrate. To test this, we determined the structure of Fbp1 alone by X-ray crystallography, which along with SEC-MALS confirmed its tetrameric assembly (Figures 2C and S1B). We then solved the structure of Fbp1 bound to the GID E3 by cryo EM, which led to several conclusions (Figure 2C). First, Fbp1 was readily docked in the extra density at the center of the large oval in the Fbp1-bound GID E3 assembly. Second, two of Fbp1’s edges approach the Gid4 subunits within each GID^SR4^ docked on opposite sides of the oval. Third, the density attributed to Gid7 does not directly contact Fbp1, but rather is localized at the vertices between two GID^SR4^ complexes. Thus, Gid7 activates GID E3 activity toward Fbp1 indirectly, by driving supramolecular assembly.

The resultant GID assembly resembles an organometallic supramolecular chelate, wherein multiple giant organic molecules capture a much smaller ligand through multiple discrete points of contact. Thus, we term the giant oval complex “Chelator-GID^SR4^”, based on its supramolecular assembly, through which two GID^SR4^ complexes simultaneously capture degrons displayed from two protomers in the tetrameric Fbp1 substrate.

### High-resolution structures of modules within Chelator-GID^SR4^

A series of focused refinements enabled building atomic models of three functionally distinct modules comprising Chelator-GID^SR4^ (Figures 3A, S1D, S2A and S3): (1) A substrate receptor scaffolding module (SRS), which is comprised of Gid1^SRS^, Gid4, Gid5, and Gid8^SRS^ in 1:1:1:1 stoichoimetry (SRS in superscript denotes Gid1 and Gid8 protomers within the substrate receptor scaffolding module) (Figure 3A, SRS panel). The SRS is responsible for bridging the substrate receptor (here Gid4, shown in red), which recruits Fbp1 for ubiquitylation, to the other E3 ligase subunits. (2) A catalytic module (Cat), which contains Gid2 and Gid9 (Figure 3A, Cat panel). Gid2’s RING domain and Gid9’s RING-like domain cooperate to bind and activate the Ubc8∼ubiquitin intermediate, from which ubiquitin is transferred from its thioester linkage with Ubc8 to a lysine on a recruited Fbp1 substrate. (3) A previously undescribed supramolecular assembly (SA) module, which contains the remaining protomers, two of Gid7, Gid1^SA^ and Gid8^SA^ (SA in superscript denotes Gid1 and Gid8 protomers within the supramolecular assembly module) (Figure 3A, SA panel).

**Figure 3.**
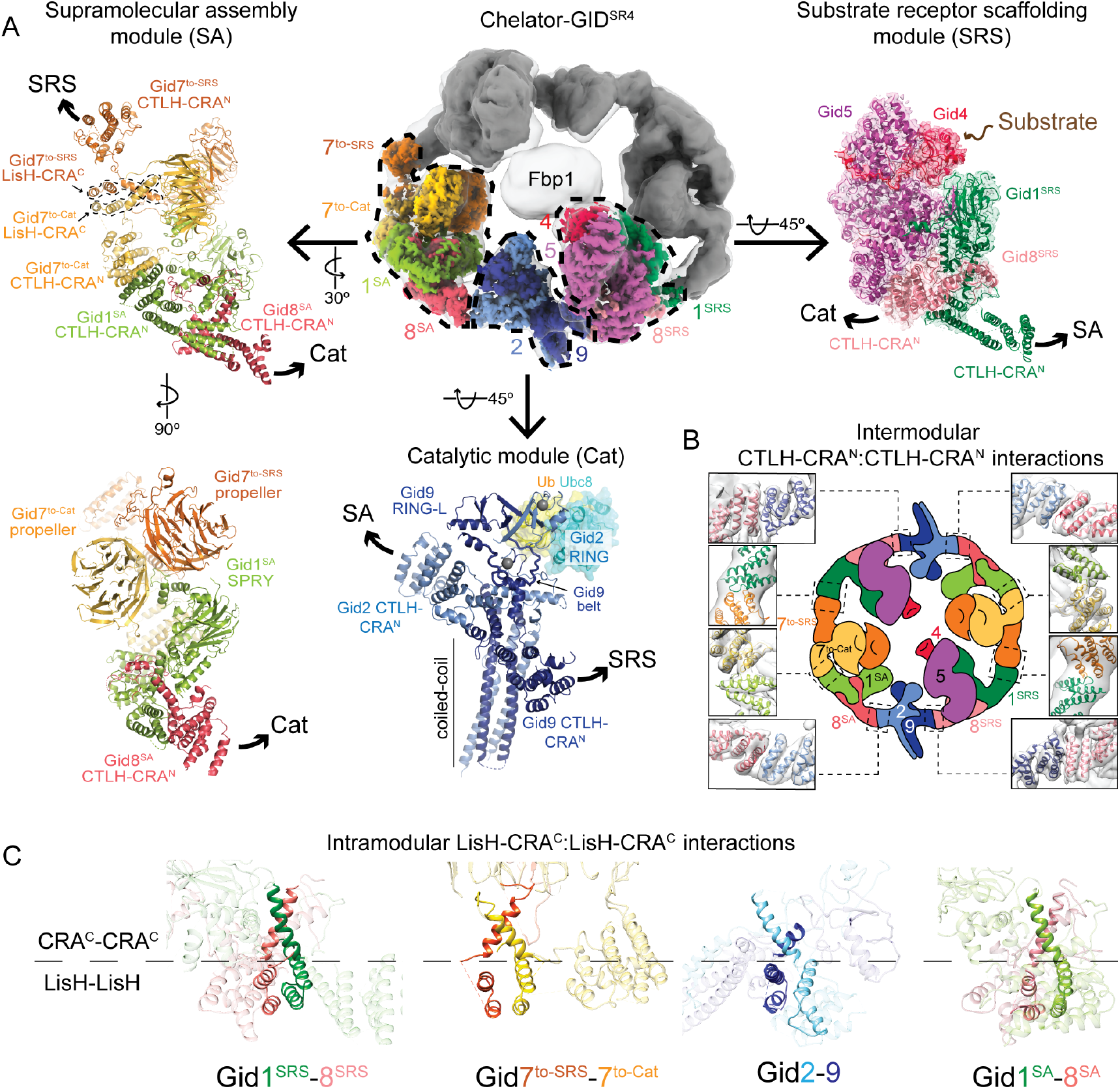
High-resolution details of Chelator-GID^SR4^ modular assembly (See also Figures S3, S4) A. Focused refined maps of substrate receptor scaffolding (SRS), catalytic (Cat) and supramolecular assembly (SA) modules are shown colored according to subunit identity, fit in half of the overall map of Fbp1-bound Chelator-GID^SR4^ (top middle). GID^SR4^ structure (6SWY.PDB) fits the SRS module (Gid1^SRS^ -dark green, Gid8^SRS^ -salmon, Gid5 -purple, Gid4 -red). Brown arrow points to Gid4’s substrate binding site (top right). Cat module comprises Gid2 (sky blue) and Gid9 (navy). Zinc ions are shown as gray spheres. Ubc8∼Ub was modelled by aligning Gid2 RING with an E2∼Ub-bound RING structure (5H7S.PDB). SA comprises Gid1^SA^ (green), Gid8^SA^ (pink) and 2 Gid7 protomers, Gid7^to-Cat^ (yellow) and Gid7^to-SRS^ (orange) facing Cat or SRS modules, respectively. Superscript refers to module for given Gid1 or Gid8 protomers. Arrows point to connected modules. B. Cartoon of Chelator-GID^SR4^, with close-ups of intermodule CTLH-CRA^N^: CTLH-CRA^N^ interactions fit into map of Chelator-GID^SR4^ (gray). C. Intramodule LisH-CRA^C^: LisH-CRA^C^ (solid ribbon) interactions in Chelator-GID^SR4^

Focused refinement of the SRS module within Chelator-GID^SR4^ yielded a map at 3.4 Å resolution, which readily accommodated the prior coordinates for this region (PDB ID: 6SWY) (Figure 3A, SRS panel). As described previously, the globular substrate-binding domain of Gid4 fits snuggly in a complementary concave surface of the elongated scaffolding subunit Gid5. This arrangement is supported by a base from Gid1^SRS^ and Gid8^SRS^, which form an intricate heterodimer involving their LisH-CTLH-CRA domains. Placing two SRS modules within Chelator-GID^SR4^ shows the degron-binding sites of the Gid4 protomers localized at the inner-most positions, facing each other toward the interior of the oval, and contacting the centrally-recruited Fbp1.

Focused refinement over the Cat module yielded a 3.8 Å resolution reconstruction (Figure 3A, Cat panel). The map quality permitted de novo building and refinement of atomic coordinates for the majority of Gid2 and Gid9, which allowed defining roles of residues known to be essential for ubiquitylation (Figure S2A and S2B). The catalytic function is mediated by a region of Gid2 that adopts an E3 ligase RING domain fold (albeit stabilized by a single zinc in the E2∼ubiquitin binding site) together with a portion of Gid9 that adopts a unique RING-Like (RING-L) structure (Braun et al., 2011; Qiao et al., 2020; Regelmann et al., 2003) (Figure S2B). Folding of the Gid2 RING depends on its incorporation into the intricately configured Gid2-Gid9 heterodimer. The Gid2 RING is embedded in an unprecedented intermolecular heart-shaped domain, surrounded by stabilizing elements from Gid9, including an intermolecular zinc-binding domain, a belt that encases roughly three-quarters of the base of Gid2’s RING, the RING-L domain that packs against the remaining side of Gid2’s RING, and the extreme C-terminus that contributes to Gid2’s RING in a manner analogous to canonical RING-RING heterodimers such as in the MDM2-MDMX complex (Linke et al., 2008; Nomura et al., 2017). Gid2 and Gid9 are further intertwined by their N-termini co-assembling in a ≈70 Å long 4-helix heterodimeric coiled-coil (Figures 3A, S2A and S2B).

Within Chelator-GID^SR4^, the two Cat modules are placed such that their heart-shaped Gid2-Gid9 RING-RING-L E3 ligase domains are positioned toward the center of the oval assembly, facing the two degron-binding Gid4 subunits. A model for the Gid2 RING-Ubc8∼ubiquitin intermediate based on published isolated RING E3-E2∼ubiquitin complexes shows the Gid2 RING domain recruiting Ubc8, and making a 3-way interface between Ubc8 and ubiquitin-linked to its active site (Figures 3A, Cat panel and S2B). Meanwhile, Gid9’s C-terminus provides residues that buttress the canonically activated RING-E2∼ubiquitin conformation (Dou et al., 2013; Plechanovova et al., 2012; Pruneda et al., 2012) (Figure S2B). This model is consistent with the effects of previously reported Gid2 and Gid9 point mutations on Fbp1 degradation (Qiao et al., 2020).

Focused refinement of the SA module within Chelator-GID^SR4^ yielded a map at 3.6 Å resolution, enabling building an atomic model for the two Gid7 protomers, Gid1^SA^ and Gid8^SA^ (Figure 3A, SA panel). The SA can be described in two halves. First, the two Gid7 protomers form an asymmetric dimer on one side of the module, and second, Gid1^SA^ and Gid8^SA^ form an interdigitated scaffold that on one side supports the Gid7 dimer and on the other extends the length of the SA module to properly connect with the Cat module.

Each Gid7 protomer consists of an N-terminal LisH-CTLH-CRA motif and an atypical β-propeller. The CTLH “domain” was originally defined as a sequence “C-terminal to LisH”, and is typically followed by a so-called CRA (CT11 and RanBPM, aka Gid1) “domain”, although crystallographic studies of other such motifs revealed that the elements do not each correspond to an individually structured domain (Martin-Arevalillo et al., 2017; Ulrich et al., 2016). Rather, LisH-CTLH-CRA motifs together form elongated helical double-sided dimerization domains. The domains are initiated with LisH and CTLH helices progressing in one direction, capped by a pair of N-terminal “CRA” helices at one end, with the remaining CRA helices reversing and traversing the length of the domain, along the way packing against CTLH helices and terminating adjacent to the LisH helices (Figure S2C). Thus, LisH sequences and the C-terminal half of the so-called CRA domain pack to form a globular LisH-CRA^C^ dimerization domain, while the CTLH and N-terminal half of the CRA sequence together form an elongated CTLH-CRA^N^ helical bundle that is a distinct dimerization domain.

Much like the LisH-CRA^C^ motifs in Gid1^SRS^ and Gid8^SRS^ and those in Gid2 and Gid9 that were previously described to mediate heterodimerization between these subunits (Qiao et al., 2020), the two Gid7 LisH-CRA^C^ motifs mediate homodimerization. The propellers from each Gid7 ensue at different relative angles, allowing for specific, asymmetric propeller-propeller interactions. The double propeller domain displays a groove that binds part of Gid1^SA^, which enables further supramolecular assembly-module-specific interactions between the CTLH-CRA^N^ domains from Gid1^SA^, a loop from Gid8^SA^, and the CTLH-CRA^N^ domain from one Gid7 protomer (Figure S2D). We term this protomer Gid7^to-cat^ because its CTLH-CRA^N^ domain faces toward the catalytic module. The remainder of the Gid1^SA^ and Gid8^SA^ subcomplex superimposes with the corresponding portion of the assembly between Gid1^SRS^ and Gid8^SRS^. At the two edges of the supramolecular assembly module, the CTLH-CRA^N^ domains from the other Gid7 protomer (Gid7^to-SRS^) and Gid8^SA^ radiate outward to directly mediate interactions with the substrate receptor scaffolding and catalytic modules, respectively.

### Supramolecular chelate assembly is supported by inter- and intra-module LisH-CTLH-CRA domain interactions

The relative arrangement of E3 ligase elements -the substrate receptor and RING domain -inside Chelator-GID^SR4^ depends on formation of the exterior oval band that holds the chelate assembly together. The oval is established by two types of inter-subunit interactions: those within but at the outer edge of each module, and those mediating inter-module connections (Figures 3B and 3C). The two types of interactions are interdependent, through a daisy-chain-like arrangement of LisH-CTLH-CRA domains. Inter-subunit interactions within the modules involve the LisH-CRA^C^ elements, with helices from each subunit wrapping around each other. As described above for the SA module, the LisH-CRA^C^ domains of the two Gid7 protomers homodimerize in a manner similar to the heterodimeric LisH-CRA^C^ domain assemblies between Gid1 and Gid8, and Gid2 and Gid9 within modules (Figure 3C).

The modules are connected to each other in Chelator-GID^SR4^ by heterotypic dimerization of CTLH-CRA^N^ domains at the edges of each module, projected outwardly in both directions (Figure 3B). Unlike inter-subunit interactions within the modules, the CTLH-CRA^N^ domains between modules bind in a side-by-side manner. The substrate receptor scaffolding and catalytic modules are adjoined by interactions between CTLH-CRA^N^ domains of Gid8^SRS^ and Gid9, forming the GID^SR4^ assembly. The catalytic and supramolecular assembly modules are bridged by interactions between CTLH-CRA^N^ domains of Gid2 and Gid8^SA^. Notably, Gid2’s CTLH-CRA^N^ domain also packs against Gid9’s RING-L domain (Figures 3A and S2B), which may explain how formation of the Chelator-GID^SR4^ assembly impacts intrinsic ubiquitin transferase activity (Figure 1C). The oval structure also depends on adjoining the substrate receptor scaffold and supramolecular assembly modules, through interactions between CTLH-CRA^N^ domains of Gid1^SRS^ and Gid7^to-SRS^. Despite the similarity of the inter-module interactions at a secondary structural level, specificity is dictated by contacts between domains ensuring formation of the Chelator-GID^SR4^ assembly.

The extended helical nature of the CTLH-CRA^N^ domains appears to facilitate subtle bending and twisting. Indeed, subtly different conformations for the exterior oval are observed in different classes from the EM data on Chelator-GID^SR4^. Intriguingly, comparing the major classes of Chelator-GID^SR4^ alone or bound to Fbp1 showed relative repositioning of the substrate receptor scaffolding module toward the center of the oval to bind substrate, resembling a venus flytrap capturing its prey (Figure 4A).

**Figure 4.**
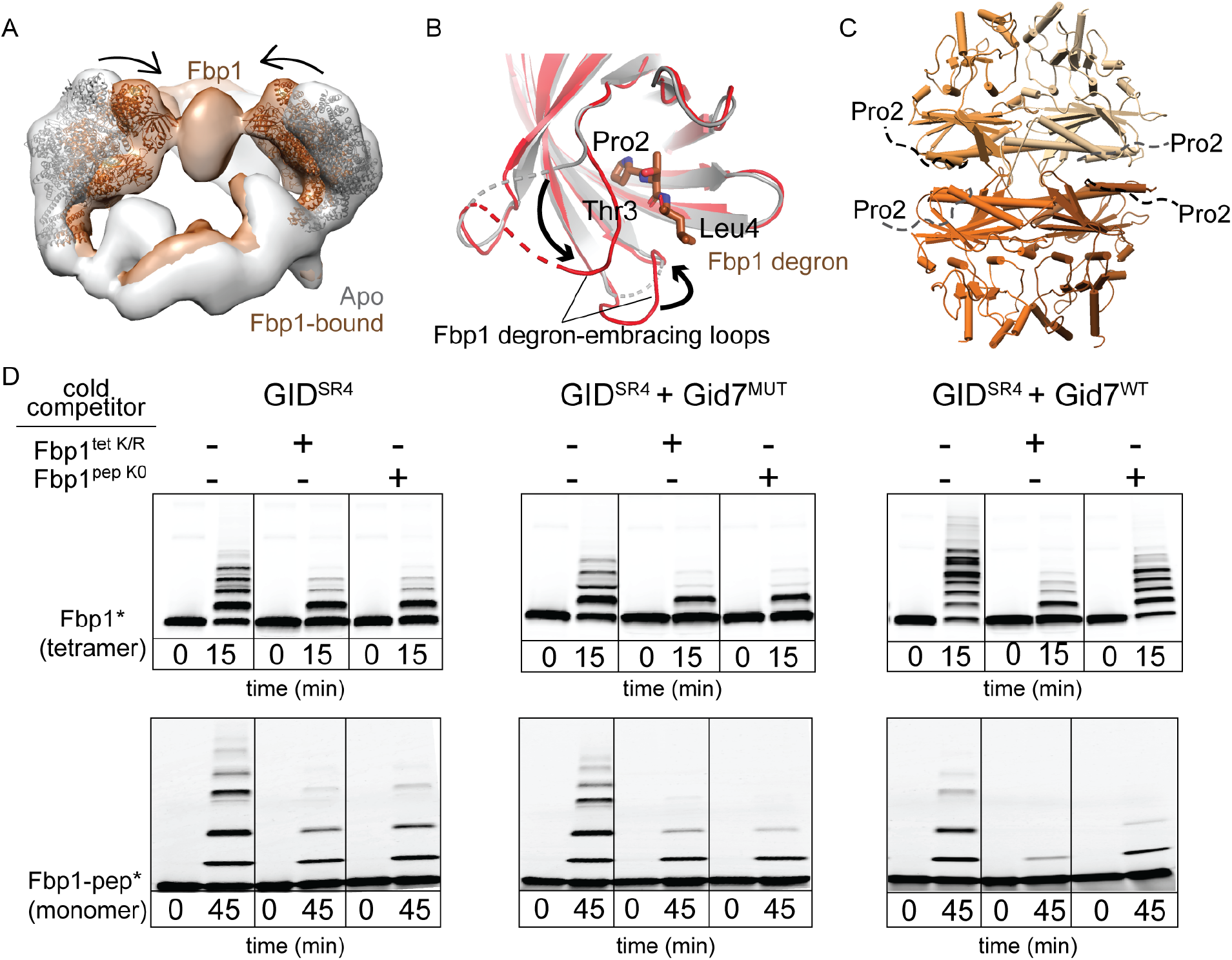
Chelator-GID^SR4^ assembly specifies multivalent binding for tetrameric Fbp1 substrate (See also Figure S4) A. Superimposed maps of substrate-free (gray) and Fbp1-bound Chelator-GID^SR4^ (brown) show relative inward movement of SRS modules (ribbon) upon substrate recruitment. B. Conformational differences between Gid4 in GID^SR4^ (6SWY.PDB, gray) and Fbp1-bound Chelator-GID^SR4^ (red). The first three residues of Fbp1 degron bound to Gid4 are shown as sticks. C. Crystal structure of Fbp1 tetramer, with N-terminal region (residues 2-19) including the degron not visible in electron density depicted as dotted lines. Fbp1 protomers shown in varying brown shades. D. Competitive *in vitro* ubiquitylation assays probing multivalent E3-substrate interactions. Chelator-GID^SR4^ has two substrate binding sites and two catalytic centers while two other E3 assemblies (GID^SR4^ or GID^SR4^ + Gid7^MUT^ lacking LisH-CTLH-CRA motif, Δ1-285) have only one substrate binding site and one catalytic center. Substrates are either oligomeric (tetrameric Fbp1) or monomeric (a peptide harboring a single acceptor Lys, Fbp1-pep) and fluorescently-labeled at the C-terminus (denoted with *). Competitors are either oligomeric (tetrameric Fbp1^tet K/R^, with preferred target lysines mutated to arginines) or monomeric (lysine-less peptide, Fbp1^pep K0^).

### Chelator-GID^SR4^ assembly mediates avid recruitment of tetrameric substrate Fbp1

To understand how the Chelator-GID^SR4^ supramolecular assembly determines regulation, we sought to generate a structural model of an active ubiquitylation complex. For an individual GID^SR4^ E3, the substrate receptor scaffold and the catalytic modules together resemble a clamp. Gid4’s degron binding site and Gid2’s RING domain correspond to the jaws that recruit and juxtapose the substrate degron and the Ubc8∼ubiquitin intermediate, respectively (Qiao et al., 2020). Thus, making a model in the context of the chelate assembly would require determining which portions of Fbp1 align with the Gid4s, and which align with the ubiquitylation active sites.

An individual Fbp1 degron motif was visually captured by Gid4 in the locally-refined map of a substrate receptor scaffolding module within Chelator-GID^SR4^ (Figure S3B). The interaction is anchored by Fbp1’s N-terminal Pro, in agreement with prior studies showing Fbp1 is a Pro/N-degron substrate, and much like prior crystal structures of a peptide bound to human Gid4 alone (Chen et al., 2017; Dong et al., 2018; Hämmerle et al., 1998). Comparing the structure presented here to the prior structure of GID^SR4^ without substrate bound showed a remodelling of several Gid4 loops, which embrace the three N-terminal residues (Pro-2, Thr-3, Leu-4) of an Fbp1 protomer (Figure 4B).

In the absence of binding Gid4, Fbp1’s N-terminal degrons were not visible in the crystal structure, suggesting that they are disordered (Figure 4C). The first visible amino acid, residue 19, from each protomer is located near the center of the disk-like Fbp1 structure. This arrangement could enable two potential degrons, one from either side, to simultaneously ensnare a Gid4 substrate receptor within Chelator-GID^SR4^. If this was the case, Chelator-GID^SR4^ would avidly bind to an Fbp1 tetramer, but not to a monomeric degron. In contrast, GID^SR4^ alone should bind equally well to degrons displayed as a monomer or from the tetrameric Fbp1 scaffold. Although we could not obtain sufficient Chelator-GID^SR4^ to quantify its interactions, we could obtain insights through competitive ubiquitylation assays (Figure 4D). Unlabeled monomeric (an isolated Fbp1 degron peptide) and oligomeric (unlabeled Fbp1) inhibitors had comparable impact on ubiquitylation of fluorescent Fbp1 by GID^SR4^, or GID^SR4^ mixed with a Gid7 mutant that does not support supramolecular assembly. However, the unlabeled Fbp1 tetramer was strikingly more effective at impeding the activity of Chelator-GID^SR4^ toward fluorescent Fbp1 than the unlabelled monomeric inhibitor. Notably, the same inhibitory trends were observed for ubiquitylation of a fluorescent monomeric peptide substrate, confirming that the Fbp1 oligomer complements the Chelator GID assembly (Figure 4D, bottom panel). These data are consistent with a model whereby avid Fbp1 recruitment to Chelator-GID^SR4^ depends on the supramolecular assemblies of both the E3 ligase and its substrate.

### Chelator-GID^SR4^ assembly establishes dual site-specific ubiquitin targeting

We mapped the regions of Fbp1 engaging the ubiquitylation catalytic centers – i.e. the thioester linkage between ubiquitin’s C-terminus and the E2 Ubc8 – by identifying its preferred sites for receiving ubiquitin. By locating di-Gly sites with mass spectrometry, we identified preferential ubiquitylation of two pairs of neighboring lysines -K32/K35 and K280/K281 -out of 18 potential target lysines on the surface of Fbp1 (Figure S4). The importance of these lysines was confirmed mutationally. Mutating the four lysines nearly eliminated Fbp1 polyubiquitylation in our *in vitro* assay, with a lesser effect of mutating either pair (K32/K35 or K280/K281) (Figure 5A, top panel). Use of K0 ubiquitin that abolishes ubiquitin chain formation showed that Fbp1 protomers are modified at up to two sites during the time-course of the experiment (Figure 5A, bottom panel). Eliminating either pair of lysines reduced this to monoubiquitylation. The results suggest that either of the two regions can be ubiquitylated independently of the other, but for a given protomer, ubiquitylation is restricted to one lysine within each of the two lysine pairs. Testing effects of the mutations on Fbp1 degradation confirmed the importance of these lysines *in vivo*, with substantial stabilization even upon mutating only the K32/K35 lysine pair (Figure 5B).

**Figure 5.**
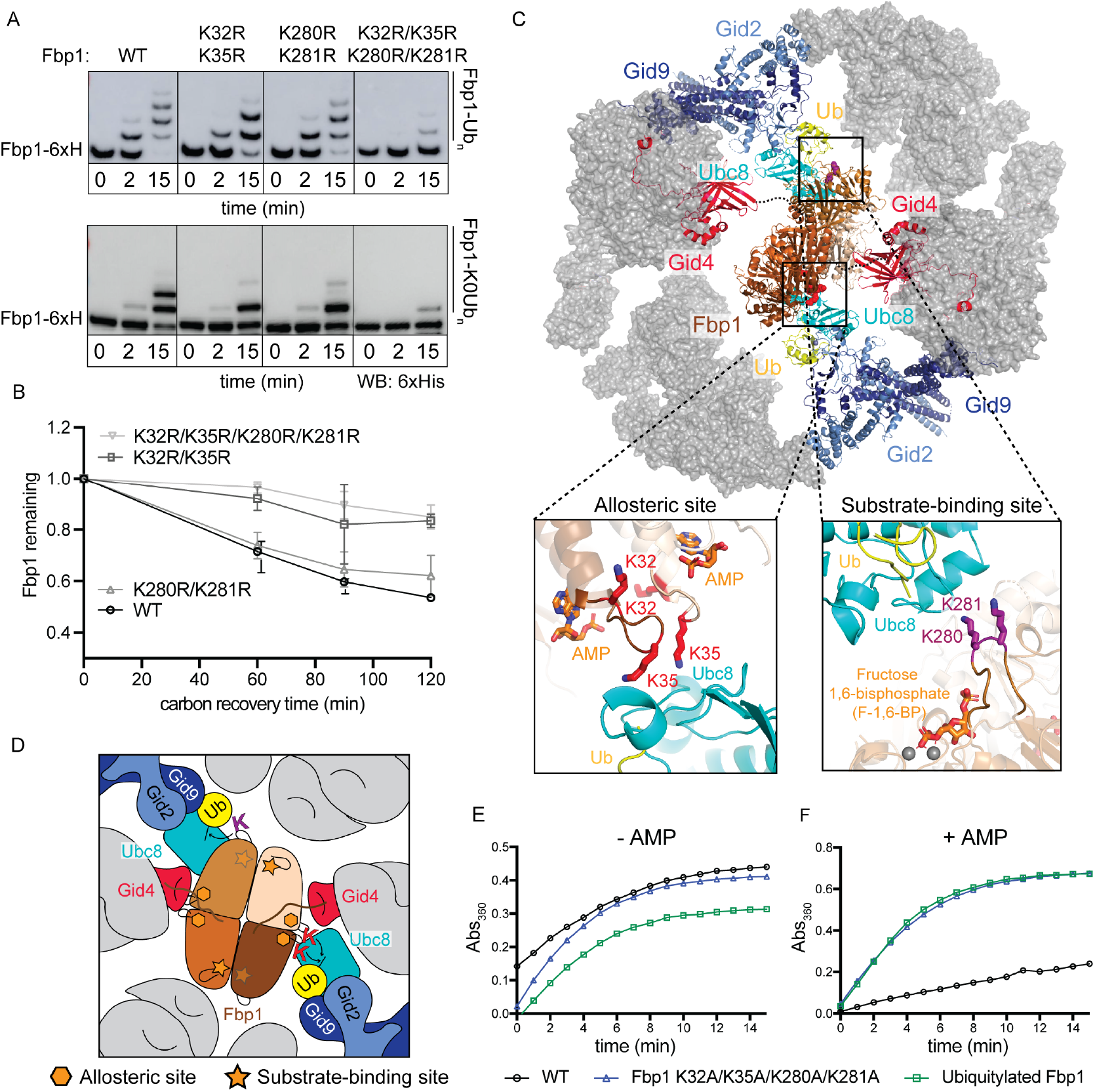
Chelator-GID^SR4^ configures simultaneous targeting of distinct lysine clusters in metabolic regulatory regions of the Fbp1 tetramer (See also Figure S5) A. *In vitro* ubiquitylation of Fbp1-6xHis, detected by anti-His immunoblot, with WT (top) or K0 (bottom) Ub, testing effects of mutating the major Fbp1 Ub-targeted lysines identified by mass spectrometry. B. Glucose-induced degradation *in vivo* of exogenously expressed WT or mutant versions of Fbp1. Substrate levels were quantified as ratio of substrate detected relative to level after the switch from carbon starvation to carbon recovery conditions. Points represent mean, error bars represent SD (n=3). C. Structural model of Chelator-GID^SR4^-mediated ubiquitylation of Fbp1. Ubc8∼Ub was modelled by aligning a homologous RING-E2∼Ub structure (5H7S.PDB) on Gid2 RING. Dotted lines indicate disordered Fbp1 N-termini. Close-ups show major Fbp1 ubiquitylation sites near substrate (F-1,6-BP) and allosteric AMP binding sides modeled from structures with human Fbp1 (5ZWK.PDB, 5ET6.PDB). D. Structure-based cartoon of Fbp1 ubiquitylation as shown in C. Stars and hexagons represent substrate-binding and the allosteric sites in Fbp1, respectively. E. *In vitro* Fbpase activity of purified WT, polyubiquitylated and mutant Fbp1 (K32A/ K35A/K280A/K281A). F. Fbpase activity assay as in E, testing responses of purified WT, polyubiquitylated and mutant Fbp1 (K32A/K35A/K280A/K281A) to allosteric inhibitor AMP.

On this basis, we modeled Fbp1 ubiquitylation as follows (Figures 5C and 5D). Fbp1 was first anchored via two degrons, one on each side binding a Gid4 in the Chelator-GID^SR4^ assembly and Ubc8∼Ub was modeled on the two Gid2 RING domains based on homology to another RING-E2∼ubiquitin assembly (Nayak and Sivaraman, 2018). Fbp1 was subjected to constrained rotation to localize the K32/K35 region of one protomer adjacent to one active site. This led to two striking observations. First, the K32/K35 regions of two pairs of protomers are adjacent to each other, allowing similar positioning of Fbp1 to ubiquitylate these sites within two protomers. Second, and unexpectedly, when a K32/K35 region is aligned with one active site, the K280/K281 region of a different Fbp1 protomer is simultaneously situated in the other active site in Chelator-GID^SR4^. Thus, the Chelator-GID^SR4^ supramolecular assembly complements the oligomeric structure of Fbp1 – enabling simultaneous capture of two Pro/N degrons, and simultaneous ubiquitylation of multiple protomers within the Fbp1 tetramer.

Because it seemed Chelator-GID^SR4^ evolved to simultaneously strike the K32/K35 and K280/K281 loops, we inspected the Fbp1 structure for potential functional importance of these regions. Intriguingly, the K32/K35 residues reside at one inter-protomer interface, in a loop abutting the allosteric site that regulates Fbp1 activity by binding the non-competitive inhibitor, AMP (Ke et al., 1990b). Meanwhile, K280/K281 are located adjacent to another inter-protomer interface in the Fbp1 tetramer -at the tip of a loop involved in binding the substrate fructose-1,6-bisphosphate (Ke et al., 1990a) (Figures 5C and 5D). We thus examined effects of Fbp1 ubiquitylation by Chelator-GID^SR4^ on its activity. A K32/K35/K280/K281 mutant, and a ubiquitylated version of Fbp1 show Fbpase activity in our assay (Figure 5E). However, allosteric modulation by AMP was substantially impaired in both cases (Figure 5F). Thus, Chelator-GID^SR4^ targets sites related to Fbp1’s metabolic function.

### Structural and mechanistic parallels in human CTLH E3

To determine if structural principles governing activity of the yeast GID E3 are conserved in higher eukaryotes, we studied the human CTLH complex, whose subunits mirror those of Chelator-GID^SR4^ (Figure 6A).

**Figure 6.**
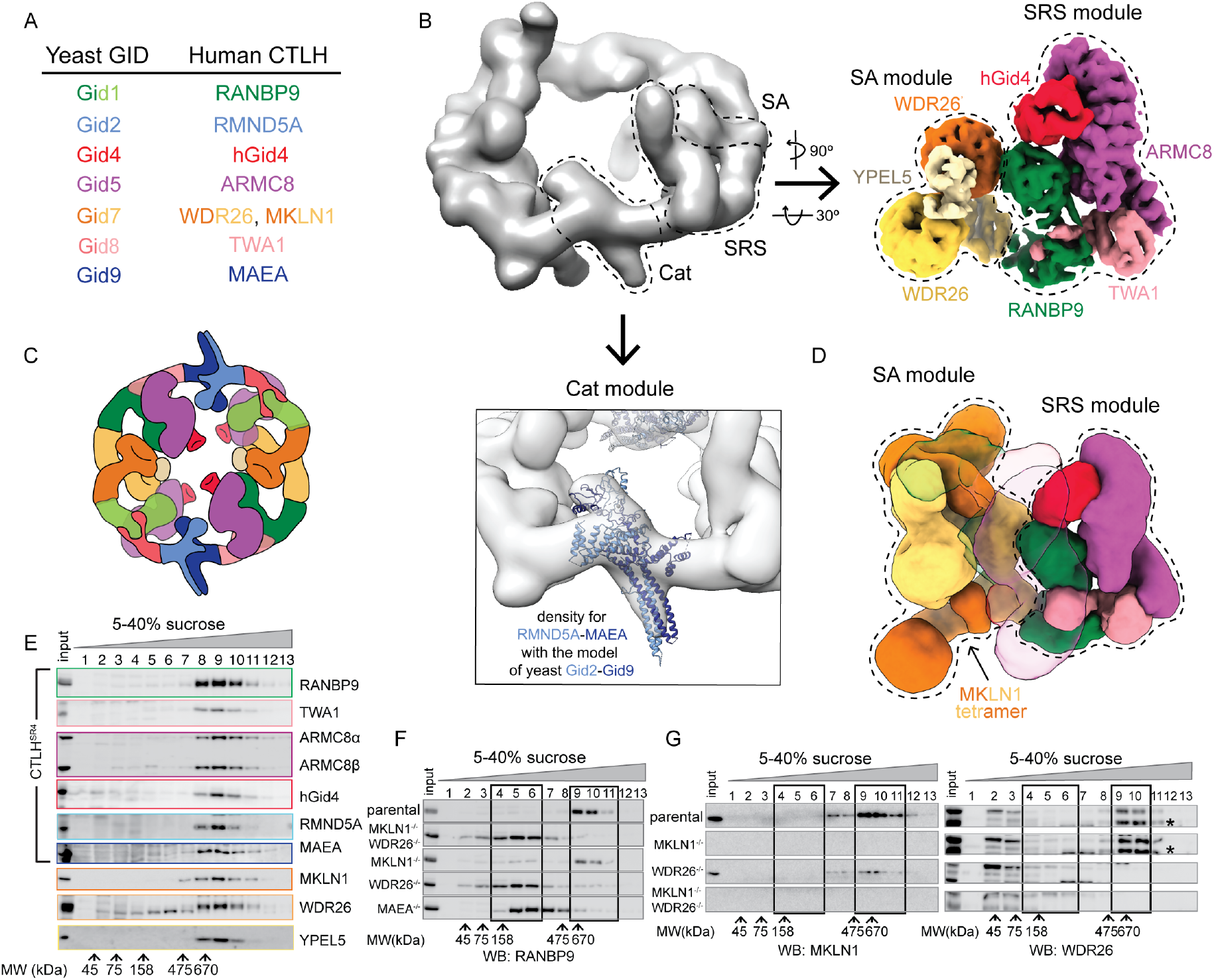
Higher order assemblies of human CTLH E3 (See also Figures S6, S7, Table S1) A. Color-coded guide to yeast GID subunits and their human orthologues in CTLH complex. Two colors indicate multiple protomers of subunit. B. Cryo EM maps of CTLH assemblies containing Cat (RMND5A-MAEA), SRS (RANBP9-TWA1-ARMC8 alone or bound to hGid4), and/or SA (WDR26 with or without YPEL5) modules, as indicated. Subunits are colored according to guide in A. Top left, low resolution map of WDR26-mediated supramolecular assembly of CTLH (RANBP9-TWA1-ARMC8-MAEA-RMND5A-WDR26). Right, 6.5 Å map of human CTLH SRS module (RANBP9-TWA1-ARMC8-hGid4) subcomplex with an SA module comprising WDR26-YPEL5. Lower panel shows yeast Gid2-Gid9 structure in corresponding CTLH Cat module. C. Cartoon colored as in A representing CTLH oval assembly wherein SA module is WDR26-YPEL5 dimer. D. 10.4 Å resolution map of human CTLH SRS module with MKLN1 as SA module. 2^nd^ copy of SRS module in subcomplex is transparent. E. Immunoblots of fractions from sucrose gradients of K562 cell lysates, probed with indicated antibodies. F. Immunoblots probing for core CTLH subunit (RANBP9) in fractions from sucrose gradients of lysates from parental K562 and WDR26^-/-^/MKLN1^-/-^, MKLN1^-/-^, WDR26^-/-^ and MAEA^-/-^ knockout cells. Black boxes delineate high and low MW peak fractions. G. As in F, but probed as indicated with anti-MKLN1 or -WDR26 antibodies. * -WDR26 band.

We first reconstituted a recombinant complex that we term “CTLH^SR4^”, which parallels yeast GID^SR4^. A low resolution cryo EM envelope showed that the corresponding human subunits form SRS (hGid4-ARMC8-RANBP9-TWA1) and Cat (RMND5A-MAEA) modules (Figure S5A). As for yeast GID^SR4^ (Qiao et al., 2020), the CTLH^SR4^ Cat module is relatively poorly resolved, but the coordinates for the yeast Gid2-Gid9 subcomplex derived from Chelator-GID^SR4^ readily fit in the density. A 3.2 Å map obtained by focused refinement enabled building atomic coordinates for the human SRS module, which superimposes with its yeast counterpart (Figures S5B, S6).

We tested if the structural conservation extended to enzymatic mechanism. Because Pro/N-end degron targets of the CTLH E3 remain unknown, we generated a model peptide substrate: an N-terminal PGLW sequence, previously reported to optimally bind hGid4 (Dong et al., 2020; Dong et al., 2018), connected via a flexible linker to a C-terminal target lysine. With this peptide substrate in-hand, we tested effects of structure-based point mutations on ubiquitylation. The hGid4 residues mediating its incorporation into CTLH^SR4^, and RMND5A and MAEA residues that activate UBE2H∼Ub, are crucial for peptide substrate ubiquitylation (Figures S5C-H). Moreover, as with GID^SR4^ (Qiao et al., 2020), only K48, out of all Ub lysines was sufficient to support polyUb chain formation by CTLH^SR4^, albeit to a substantially lesser degree than WT Ub (Figure S5I). Thus, it seems the human CTLH core module parallels that in yeast GID assemblies.

We examined by cryo EM if the human Gid7 orthologues, WDR26 and MKLN1, have capacity for supramolecular assembly. We obtained reconstructions for two subcomplexes containing WDR26. Coexpressing WDR26 with scaffolding and catalytic subunits (ARMC8-RANBP9-TWA1-RMND5A-MAEA) yielded a complex broadly resembling Chelator-GID^SR4^ in that it forms a hollow oval of similar dimensions (Figures 6B and 6C). Docking structures of human and yeast subcomplexes into the density showed, however, that a WDR26 dimer itself is the supramolecular assembly module. WDR26 binds directly to RANBP9-TWA1 in the scaffold, without duplicates of these subunits corresponding to yeast Gid1^SA^-Gid8^SA^. The distinct WDR26-dependent supramolecular assembly places four – not two – ARMC8 subunits poised to each bind a hGid4 to capture substrate degrons in the CTLH oval.

The distinctive arrangement of SA and SRS modules was preserved in a 6 Å resolution map of WDR26, RANBP9, TWA1, ARMC8, hGid4 and the poorly understood CTLH subunit, YPEL5 (Figure 6B). Interestingly, YPEL5 binds at the junction of the two protomers in the WDR26 double propeller domain.

A low resolution map showed yet another supramolecular assembly for other human Gid7 ortholog, MKLN1, bound to the CTLH SRS module (Figure 6D). Like WDR26, MKLN1 binds directly to RANBP9-TWA1 in the scaffold, without intervening duplicates of these subunits. However, in accordance with previous studies (Delto et al., 2015; Kim et al., 2014), MKLN1 forms a tetramer. Four MKLN1 protomers bind between two CTLH SRS modules, demonstrating potential for even higher-order CTLH complex assemblies.

We confirmed roles of WDR26 and MKLN1 in human CTLH complex assembly, by sedimentation analyses of lysates from K562 cells, or lines in which the human Gid7 orthologs were deleted. Immunoblotting of fractions from sucrose density gradients of parental K562 cell lysates showed comigration of CTLH subunits, corresponding to a complex of molecular weight greater than that predicted for a uniformly stoichiometric assembly (600-800 kDa according to standards) (Figure 6E). However, probing migration of core subunit RANBP9 as a marker for the CTLH complex showed that the assembly changes markedly -towards fractions of 150-350 kDa -in CRISPR/Cas9 genome-edited lines lacking WDR26, MKLN1, or both, or the Cat module subunit MAEA (Figures 6F and S5J). Interestingly, migration of WDR26 and MKLN1 in higher molecular weight fractions is not interdependent (Figure 6G), possibly indicating that each Gid7 ortholog can reside in distinct CTLH assemblies. Much of the total CTLH population shifted to lower molecular weight fractions upon deletion of WDR26, with lesser effect of deleting MKLN1. This may suggest a greater proportion of the CTLH complex in these cells depends on WDR26 for supramolecular assembly, perhaps due to a higher relative concentration of WDR26, or factors differentially regulating WDR26 or MKLN1 assembly into CTLH complexes.

Overall, the results suggest that CTLH E3 assemblies contain SRS, Cat and SA modules with features resembling those in Chelator-GID^SR4^. Moreover, differences in structural configurations of complexes containing MKLN1 or WDR26 offer prospects that CTLH may adopt a variety of supramolecular E3 assemblies, which could further impart distinct functionalities.

## DISCUSSION

Here we discovered multipronged substrate targeting by an E3 ligase chelate supramolecular assembly tailored to the oligomeric quaternary structure of its metabolic enzyme substrate. In the absence of chelate assembly, GID^SR4^ is a competent E3 ligase that can bind a substrate degron, activate the intrinsic reactivity of its E2 partner -the Ubc8∼Ub intermediate - and promote Ub transfer from Ubc8 to a recruited substrate (Qiao et al., 2020). GID^SR4^ is also competent *in vitro* insofar as Gid7 is not required for glucose - and GID-dependent degradation of Mdh2 (Qiao et al., 2020). Instead of binding directly to its specified substrate Fbp1, Gid7 alters the GID assembly (Figures 1, 2, 3).

While other E3s have been reported to self-assemble (Balaji and Hoppe, 2020), this typically is achieved by catalytic or substrate receptor subunits, for example the dimeric RING domains of single subunit E3s or dimeric F-box and BTB substrate receptors in multisubunit cullin-RING ligases (Dou et al., 2012; McMahon et al., 2006; Ogura et al., 2010; Plechanovova et al., 2012; Welcker et al., 2013; Zhuang et al., 2009). Substrate-bound multivalent E3s can undergo liquid-liquid phase-separation (Bouchard et al., 2018). However, the transformation into Chelator-GID^SR4^ represents a distinctive, extreme, and specific adjustment of an E3 ligase architecture (Figures 2 and 3).

Resembling an organometallic chelate interacting with its central ligand, Chelator-GID^SR4^’s multiple distinct points of contact with Fbp1 not only include the degron-binding sites from two opposing Gid4 substrate receptors, but also the ubiquitylation active sites from Ubc8∼Ub intermediates activated by two opposing Gid2-Gid9 catalytic domains (Figures 4, 5 and 7). Relative to the monodentate GID^SR4^, the Chelator-GID^SR4^ assembly enables more molecules within the Fbp1 tetramer to be simultaneously ubiquitylated, thereby increasing Ub density on a given Fbp1 tetramer (Figure 1B). Interestingly, there is not a 1:1 correspondence between number of degron binding sites in Chelator-GID^SR4^ and number of degrons in Fbp1. The Fbp1 tetramer has four potential degrons exposed, two on each side, both seemingly poised to capture one central-facing Gid4 in Chelator-GID^SR4^ (Figure 4C). An excess number of degrons is reminiscent of substrates recruited to the cullin-RING ligase receptor Cdc4, whose single binding site can continually and dynamically sample multiple degrons (Mittag et al., 2008). For Chelator-GID^SR4^-bound Fbp1, we speculate that the arrangement of degrons allows their rapid interchange. This could potentially mediate switching between the protomers positioned adjacent to the active sites.

**Figure 7.**
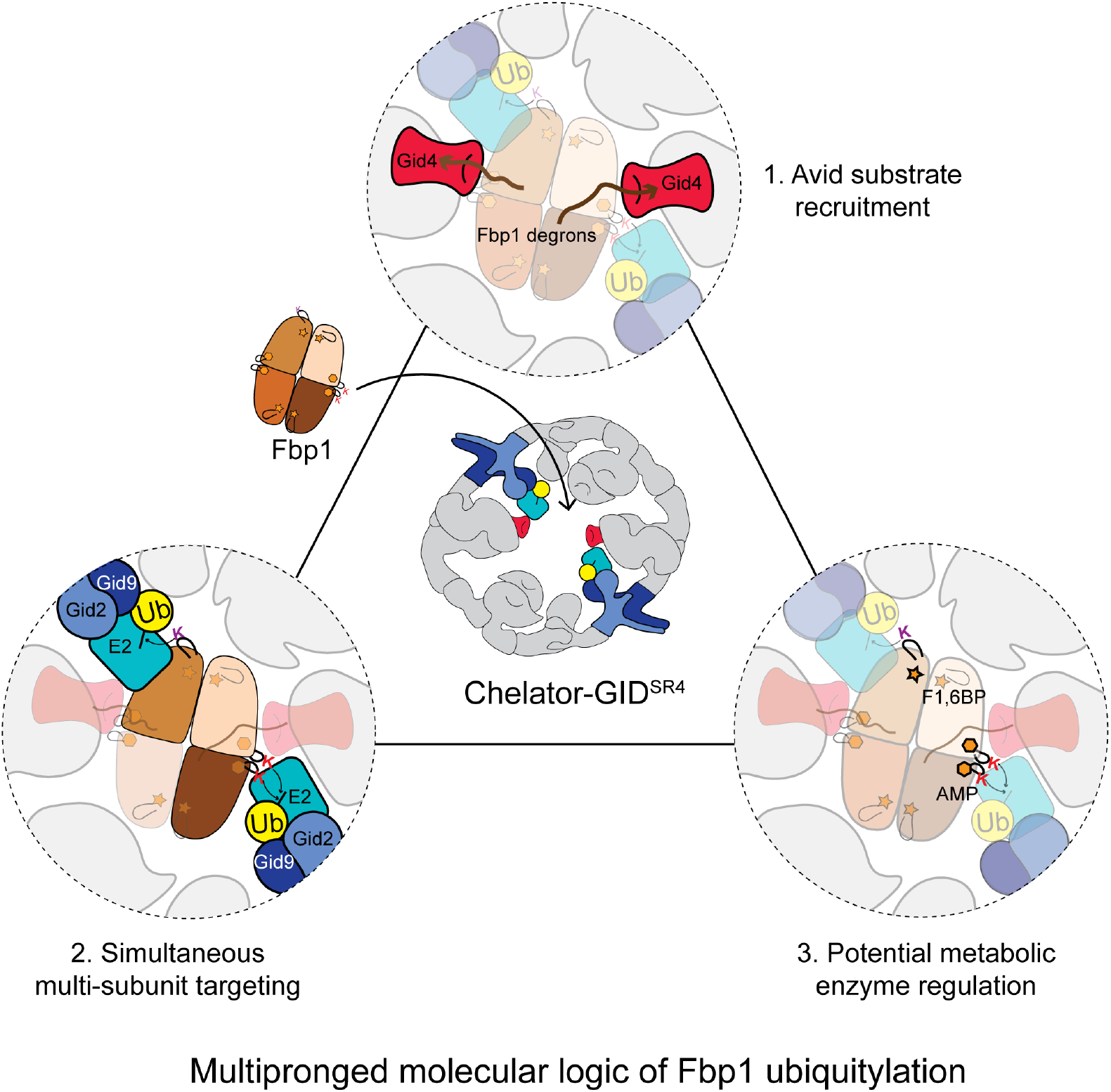
Molecular logic of multipronged Ub targeting of Fbp1 by Chelator-GID^SR4^. Supramolecular chelate assembly specifies oligomeric metabolic enzyme targeting: 1) Opposing Gid4 subunits avidly bind multiple degrons of tetrameric Fbp1; 2) Opposing RING-E2∼Ub active sites simultaneously target specific lysine clusters; 3) Targeted lysines map to metabolically-important regions of oligomeric substrate.

The human CTLH E3 complex displays striking parallels to Chelator-GID^SR4^, albeit with interesting twists. In particular, the different Gid7 orthologs form distinct supramolecular assemblies (Figure 6). We speculate that the unique assemblies define distinct functions, as implied by varying phenotypic alterations upon their individual mutation (Bauer et al., 2018; Nassan et al., 2017; Skraban et al., 2017; Zhen et al., 2020).

Taken together with previous data (Lampert et al., 2018; Qiao et al., 2020), it is now clear there is not a single yeast GID or human CTLH complex. Rather, GID and CTLH provide examples of responsive systems of multiprotein assemblies -with an active E3 core that can be elaborated by supramolecular assembly. While the function of one such assembly was shown here, the variations revealed by human Gid7 orthologs suggest that they and presumably other subunits also co-configure substrate binding and ubiquitylation active sites in accordance with molecular organization and quarternary structure of particular substrates. The Chelator model presented here demonstrates how GID (and presumably CTLH) utilizes an elegant molecular logic: the response to glucose availability converges on numerous aspects of its substrate’s structure and function to achieve precise physiological regulation (Figure 7).

## OPEN QUESTIONS

Chelator-GID^SR4^ is remarkably specific in ubiquitylating particular Fbp1 lysines within metabolic regulatory regions. However, physiologic roles of Fbp1 ubiquitylation impairing allosteric regulation and metabolic function are unknown. Future studies will be required to determine how metabolic flux is coupled with GID-dependent ubiquitylation during the termination of gluconeogenesis.

Although Chelator-GID^SR4^ is active toward Mdh2, it is unclear why this oligomeric substrate is less dependent than Fbp1 on Gid7-mediated supramolecular assembly. One speculative possibility could be that any potential advantage of avid binding is offset by accessibility of numerous ubiquitylation sites to GID^SR4^. Future studies will be required to understand how other GID E3 substrates including the Gid4 substrate receptor itself, are recognized and ubiquitylated (Hämmerle et al., 1998; Karayel et al., 2020; Menssen et al., 2018).

Finally, although the discovery of the Chelator configuration provides a basis for understanding GID higher order assembly, what other assemblies or sub-assemblies may form, and their functions remain unknown. Clearly, other arrangements are observed for human CTLH complexes with WDR26. MKLN1 forms yet an even higher-order assembly with the human SRS module. Meanwhile, some yeast GID assemblies migrate in the void volume by size-exclusion chromatography (Figure S1A). Moreover, mechanistic roles of additional subunits, including YPEL5 (Figure 6) remain unknown (Lampert et al., 2018). We thus await future studies revealing functions of other variations on GID and CTLH assemblies.

## ACKNOWLEDGEMENTS

We thank A. Varshavsky for plasmids for promoter reference assays in yeast; S. Übel and S. Pettera from MPIB biochemistry core facility; D. Bollschweiler and T. Schäfer from MPIB Cryo-EM facility; J. Basquin, K. Valer-Saldana, S. Pleyer from MPIB crystallography facility, Swiss Light Source for crystal data collection, J. Frye for initial cloning of hGID subunits, M. Yamaguchi for cloning yeast Gid7 constructs, K. Baek, B. Bräuning for helpful suggestions and all other members of Schulman lab for their support. BAS has received funding from the European Research Council (ERC) under the European Union’s Horizon 2020 research and innovation programme (grant agreement No 789016-NEDD8Activate), and the Leibniz Prize from the Deutsche Forschungsgemeinschaft (DFG, German Research Foundation -SCHU 3196/1-1). BAS and MM are supported by the Max Planck Society.

## AUTHOR CONTRIBUTIONS

Conceptualization, D.S., J.C., A.F.A. and B.A.S.; Methodology, D.S., J.C., J.R.P., C.R.L., O.K., A.F.A. and B.A.S.; Investigation, D.S., J.C., S.Q., L.A.H., C.R.L., K.G., O.K., F.M.H. and A.F.A.; Resources, S.v.G. and J.R.P.; Writing – Original Draft, D.S., J.C. and B.A.S.; Writing – Review & Editing D.S., J.C., C.R.L., A.F.A and B.A.S.; Supervision, M.M., A.F.A. and B.A.S.; Funding Acquisition, M.M. and B.A.S.

## REFERENCES

Baek, K., Krist, D.T., Prabu, J.R., Hill, S., Klügel, M., Neumaier, L., von Gronau, S., Kleiger, G., and Schulman, B.A. (2020). NEDD8 nucleates a multivalent cullin–RING– UBE2D ubiquitin ligation assembly. Nature 578.

Balaji, V., and Hoppe, T. (2020). Regulation of E3 ubiquitin ligases by homotypic and heterotypic assembly. F1000Res 9.

Barford, D. (2020). Structural interconversions of the anaphase-promoting complex/cyclosome (APC/C) regulate cell cycle transitions. Curr Opin Struct Biol 61, 86–97.

Bauer, A., Jagannathan, V., Hogler, S., Richter, B., McEwan, N.A., Thomas, A., Cadieu, E., Andre, C., Hytonen, M.K., Lohi, H., et al. (2018). MKLN1 splicing defect in dogs with lethal acrodermatitis. PLoS Genet 14, e1007264.

Boldt, K., van Reeuwijk, J., Lu, Q., Koutroumpas, K., Nguyen, T.M., Texier, Y., van Beersum, S.E., Horn, N., Willer, J.R., Mans, D.A., et al. (2016). An organelle-specific protein landscape identifies novel diseases and molecular mechanisms. Nat Commun 7, 11491.

Bouchard, J.J., Otero, J.H., Scott, D.C., Szulc, E., Martin, E.W., Sabri, N., Granata, D., Marzahn, M.R., Lindorff-Larsen, K., Salvatella, X., et al. (2018). Cancer Mutations of the Tumor Suppressor SPOP Disrupt the Formation of Active, Phase-Separated Compartments. Mol Cell 72, 19–36 e18.

Braun, B., Pfirrmann, T., Menssen, R., Hofmann, K., Scheel, H., and Wolf, D.H. (2011). Gid9, a second RING finger protein contributes to the ubiquitin ligase activity of the Gid complex required for catabolite degradation. FEBS Lett 585, 3856–3861.

Brown, N.G., VanderLinden, R., Watson, E.R., Weissmann, F., Ordureau, A., Wu, K.P., Zhang, W., Yu, S., Mercredi, P.Y., Harrison, J.S., et al. (2016). Dual RING E3 Architectures Regulate Multiubiquitination and Ubiquitin Chain Elongation by APC/C. Cell 165, 1440–1453.

Cannon, K.A., Ochoa, J.M., and Yeates, T.O. (2019). High-symmetry protein assemblies: patterns and emerging applications. Curr Opin Struct Biol 55, 77–84.

Chen, S., Wu, J., Lu, Y., Ma, Y.B., Lee, B.H., Yu, Z., Ouyang, Q., Finley, D.J., Kirschner, M.W., and Mao, Y. (2016). Structural basis for dynamic regulation of the human 26S proteasome. Proc Natl Acad Sci U S A 113, 12991–12996.

Chen, S.J., Wu, X., Wadas, B., Oh, J.H., and Varshavsky, A. (2017). An N-end rule pathway that recognizes proline and destroys gluconeogenic enzymes. Science 355.

Chiang, H.L., and Schekman, R. (1991). Regulated import and degradation of a cytosolic protein in the yeast vacuole. Nature 350, 313–318.

Delto, C.F., Heisler, F.F., Kuper, J., Sander, B., Kneussel, M., and Schindelin, H. (2015). The LisH motif of muskelin is crucial for oligomerization and governs intracellular localization. Structure 23, 364–373.

Dong, C., Chen, S.J., Melnykov, A., Weirich, S., Sun, K., Jeltsch, A., Varshavsky, A., and Min, J. (2020). Recognition of nonproline N-terminal residues by the Pro/N-degron pathway. Proc Natl Acad Sci U S A 117, 14158–14167.

Dong, C., Zhang, H., Li, L., Tempel, W., Loppnau, P., and Min, J. (2018). Molecular basis of GID4-mediated recognition of degrons for the Pro/N-end rule pathway. Nat Chem Biol 14, 466–473.

Dou, H., Buetow, L., Sibbet, G.J., Cameron, K., and Huang, D.T. (2013). Essentiality of a non-RING element in priming donor ubiquitin for catalysis by a monomeric E3. Nat Struct Mol Biol 20, 982–986.

Francis, O., Han, F., and Adams, J.C. (2013). Molecular phylogeny of a RING E3 ubiquitin ligase, conserved in eukaryotic cells and dominated by homologous components, the muskelin/RanBPM/CTLH complex. PLoS One 8, e75217.

Gancedo, C. (1971). Inactivation of Fructose-1,6-Diphosphatase by Glucose in Yeast. Journal of Bacteriology 107.

Hämmerle, M., Baue, r.J., Rose, M., Szallies, A., Thumm, M., Düsterhus, S., Mecke, D., Entian, K.D., and Wolf, D.H. (1998). Proteins of Newly Isolated Mutants and the Amino-terminal Proline Are Essential for UbiquitinProteasome-catalyzed Catabolite Degradation of Fructose-1,6-bisphosphatase of Saccharomyces cerevisiae. The Journal of Biological Chemistry 273.

Han, T., Yang, C.S., Chang, K.Y., Zhang, D., Imam, F.B., and Rana, T.M. (2016). Identification of novel genes and networks governing hematopoietic stem cell development. EMBO Rep 17, 1814–1828.

Haselbach, D., Schrader, J., Lambrecht, F., Henneberg, F., Chari, A., and Stark, H. (2017). Long-range allosteric regulation of the human 26S proteasome by 20S proteasome-targeting cancer drugs. Nat Commun 8, 15578.

Huang, D., Friesen, H., and Andrews, B. (2007). Pho85, a multifunctional cyclin-dependent protein kinase in budding yeast. Mol Microbiol 66, 303–314.

Hunkeler, M., Hagmann, A., Stuttfeld, E., Chami, M., Guri, Y., Stahlberg, H., and Maier, T. (2018). Structural basis for regulation of human acetyl-CoA carboxylase. Nature 558, 470–474.

Izard, T., Aevarsson, A., Allen, M.D., Westphal, A.H., Perham, R.N., de Kok, A., and Hol, W.G. (1999). Principles of quasi-equivalence and Euclidean geometry govern the assembly of cubic and dodecahedral cores of pyruvate dehydrogenase complexes. Proc Natl Acad Sci U S A 96, 1240–1245.

Javan, G.T., Salhotra, A., Finley, S.J., and Soni, S. (2018). Erythroblast macrophage protein (Emp): Past, present, and future. Eur J Haematol 100, 3–9.

Ke, H.M., Thorpe, C.M., Seaton, B., Lipscomb, W.N., and Marcus, F. (1990a). Structure refinement of fructose-1,6-bisphosphatase and its fructose 2,6-bisphosphate complex at 2.8 A resolution. J Mol Biol 212, 513–539.

Ke, H.M., Zhang, Y.P., and Lipscomb, W.N. (1990b). Crystal structure of fructose-1,6- bisphosphatase complexed with fructose 6-phosphate, AMP, and magnesium. Proc Natl Acad Sci U S A 87, 5243–5247.

Kim, K.H., Hong, S.K., Hwang, K.Y., and Kim, E.E. (2014). Structure of mouse muskelin discoidin domain and biochemical characterization of its self-association. Acta Crystallogr D Biol Crystallogr 70, 2863–2874.

Kobayashi, N., Yang, J., Ueda, A., Suzuki, T., Tomaru, K., Takeno, M., Okuda, K., and Ishigatsubo, Y. (2007). RanBPM, Muskelin, p48EMLP, p44CTLH, and the armadillo-repeat proteins ARMC8alpha and ARMC8beta are components of the CTLH complex. Gene 396, 236–247.

Koshland, D.E., Jr. (1963a). Correlation of Structure and Function in Enzyme Action. Science 142, 1533–1541.

Koshland, D.E., Jr. (1963b). Properties of the active site of enzymes. Ann N Y Acad Sci 103, 630–642.

Lampert, F., Stafa, D., Goga, A., Soste, M.V., Gilberto, S., Olieric, N., Picotti, P., Stoffel, M., and Peter, M. (2018). The multi-subunit GID/CTLH E3 ubiquitin ligase promotes cell proliferation and targets the transcription factor Hbp1 for degradation. Elife 7.

Lander, G.C., Estrin, E., Matyskiela, M.E., Bashore, C., Nogales, E., and Martin, A. (2012). Complete subunit architecture of the proteasome regulatory particle. Nature 482, 186–191.

Linke, K., Mace, P.D., Smith, C.A., Vaux, D.L., Silke, J., and Day, C.L. (2008). Structure of the MDM2/MDMX RING domain heterodimer reveals dimerization is required for their ubiquitylation in trans. Cell Death Differ 15, 841–848.

Liu, H., and Pfirrmann, T. (2019). The Gid-complex: an emerging player in the ubiquitin ligase league. Biol Chem 400, 1429–1441.

Maitland, M.E.R., Onea, G., Chiasson, C.A., Wang, X., Ma, J., Moor, S.E., Barber, K.R., Lajoie, G.A., Shaw, G.S., and Schild-Poulter, C. (2019). The mammalian CTLH complex is an E3 ubiquitin ligase that targets its subunit muskelin for degradation. Sci Rep 9, 9864.

Martin-Arevalillo, R., Nanao, M.H., Larrieu, A., Vinos-Poyo, T., Mast, D., Galvan-Ampudia, C., Brunoud, G., Vernoux, T., Dumas, R., and Parcy, F. (2017). Structure of the Arabidopsis TOPLESS corepressor provides insight into the evolution of transcriptional repression. Proc Natl Acad Sci U S A 114, 8107–8112.

Mayhew, M., da Silva, A.C., Martin, J., Erdjument-Bromage, H., Tempst, P., and Hartl, F.U. (1996). Protein folding in the central cavity of the GroEL-GroES chaperonin complex. Nature 379, 420–426.

McMahon, M., Thomas, N., Itoh, K., Yamamoto, M., and Hayes, J.D. (2006). Dimerization of substrate adaptors can facilitate cullin-mediated ubiquitylation of proteins by a “tethering” mechanism: a two-site interaction model for the Nrf2-Keap1 complex. J Biol Chem 281, 24756–24768.

Melnykov, A., Chen, S.J., and Varshavsky, A. (2019). Gid10 as an alternative N-recognin of the Pro/N-degron pathway. Proc Natl Acad Sci U S A 116, 15914–15923.

Menssen, R., Bui, K., and Wolf, D.H. (2018). Regulation of the Gid ubiquitin ligase recognition subunit Gid4. FEBS Lett 592, 3286–3294.

Menssen, R., Schweiggert, J., Schreiner, J., Kusevic, D., Reuther, J., Braun, B., and Wolf, D.H. (2012). Exploring the topology of the Gid complex, the E3 ubiquitin ligase involved in catabolite-induced degradation of gluconeogenic enzymes. J Biol Chem 287, 25602–25614.

Mittag, T., Orlicky, S., Choy, W.Y., Tang, X., Lin, H., Sicheri, F., Kay, L.E., Tyers, M., and Forman-Kay, J.D. (2008). Dynamic equilibrium engagement of a polyvalent ligand with a single-site receptor. Proc Natl Acad Sci U S A 105, 17772–17777.

Monod, J., Changeux, J.P., and Jacob, F. (1963). Allosteric proteins and cellular control systems. J Mol Biol 6, 306–329.

Nakatsukasa, K., Okumura, F., and Kamura, T. (2015). Proteolytic regulation of metabolic enzymes by E3 ubiquitin ligase complexes: lessons from yeast. Crit Rev Biochem Mol Biol 50, 489–502.

Nassan, M., Li, Q., Croarkin, P.E., Chen, W., Colby, C.L., Veldic, M., McElroy, S.L., Jenkins, G.D., Ryu, E., Cunningham, J.M., et al. (2017). A genome wide association study suggests the association of muskelin with early onset bipolar disorder: Implications for a GABAergic epileptogenic neurogenesis model. J Affect Disord 208, 120–129.

Nayak, D., and Sivaraman, J. (2018). Structure of LNX1:Ubc13∼Ubiquitin Complex Reveals the Role of Additional Motifs for the E3 Ligase Activity of LNX1. J Mol Biol 430, 1173–1188.

Negoro, H., Matsumura, K., Matsuda, F., Shimizu, H., Hata, Y., and Ishida, H. (2020). Effects of mutations of GID protein-coding genes on malate production and enzyme expression profiles in Saccharomyces cerevisiae. Appl Microbiol Biotechnol 104, 4971–4983.

Nguyen, A.T., Prado, M.A., Schmidt, P.J., Sendamarai, A.K., Wilson-Grady, J.T., Min, M., Campagna, D.R., Tian, G., Shi, Y., Dederer, V., et al. (2017). UBE2O remodels the proteome during terminal erythroid differentiation. Science 357.

Nomura, K., Klejnot, M., Kowalczyk, D., Hock, A.K., Sibbet, G.J., Vousden, K.H., and Huang, D.T. (2017). Structural analysis of MDM2 RING separates degradation from regulation of p53 transcription activity. Nat Struct Mol Biol 24, 578–587.

Ogura, T., Tong, K.I., Mio, K., Maruyama, Y., Kurokawa, H., Sato, C., and Yamamoto, M. (2010). Keap1 is a forked-stem dimer structure with two large spheres enclosing the intervening, double glycine repeat, and C-terminal domains. Proc Natl Acad Sci U S A 107, 2842–2847.

Oh, J.H., Chen, S.J., and Varshavsky, A. (2017). A reference-based protein degradation assay without global translation inhibitors. J Biol Chem 292, 21457–21465.

Pfirrmann, T., Villavicencio-Lorini, P., Subudhi, A.K., Menssen, R., Wolf, D.H., and Hollemann, T. (2015). RMND5 from Xenopus laevis is an E3 ubiquitin-ligase and functions in early embryonic forebrain development. PLoS One 10, e0120342.

Plechanovova, A., Jaffray, E.G., Tatham, M.H., Naismith, J.H., and Hay, R.T. (2012). Structure of a RING E3 ligase and ubiquitin-loaded E2 primed for catalysis. Nature 489, 115–120.

Pruneda, J.N., Littlefield, P.J., Soss, S.E., Nordquist, K.A., Chazin, W.J., Brzovic, P.S., and Klevit, R.E. (2012). Structure of an E3:E2∼Ub complex reveals an allosteric mechanism shared among RING/U-box ligases. Mol Cell 47, 933–942.

Qiao, S., Langlois, C.R., Chrustowicz, J., Sherpa, D., Karayel, O., Hansen, F.M., Beier, V., von Gronau, S., Bollschweiler, D., Schafer, T., et al. (2020). Interconversion between Anticipatory and Active GID E3 Ubiquitin Ligase Conformations via Metabolically Driven Substrate Receptor Assembly. Mol Cell 77, 150–163 e159.

Regelmann, J., Schüle, T., Josupeit, F.S., Horak, J., Rose, M., Entian, K.D., Thumm, M., and Wolf, D.H. (2003). Catabolite Degradation of Fructose-1,6-bisphosphatase in the Yeast Saccharomyces cerevisiae: A Genome-wide Screen Identifies Eight Novel GID Genes and Indicates the Existence of Two Degradation Pathways. Molecular Biology of the Cell 14, 1652–1663.

Rosenthal, P.B., and Henderson, R. (2003). Optimal determination of particle orientation, absolute hand, and contrast loss in single-particle electron cryomicroscopy. J Mol Biol 333, 721–745.

Rusnac, D.V., and Zheng, N. (2020). Structural Biology of CRL Ubiquitin Ligases. Adv Exp Med Biol 1217, 9–31.

Salemi, L.M., Maitland, M.E.R., McTavish, C.J., and Schild-Poulter, C. (2017). Cell signalling pathway regulation by RanBPM: molecular insights and disease implications. Open Biol 7.

Santt, O., Pfirrmann, T., Braun, B., Juretschke, J., Kimmig, P., Scheel, H., Hofmann, K., Thumm, M., and Wolf, D.H. (2008). The yeast GID complex, a novel ubiquitin ligase (E3) involved in the regulation of carbohydrate metabolism. Mol Biol Cell 19, 3323–3333.

Schork, S.M., Bee, G., Thumm, M., and Wolf, D.H. (1994a). Catabolite inactivation of fructose-1,6-bisphosphatase in yeast is mediated by the proteasome. FEBS Lett 349, 270–274.

Schork, S.M., Bee, G., Thumm, M., and Wolf, D.H. (1994b). Catabolite inactivation of fructose- 1,6_bisphosphatase in yeast is mediated by the proteasome. FEBS Letters.

Schork, S.M., Bee, G., Thumm, M., and Wolf, D.H. (1994c). Site of catabolite inactivation. Nature 369, 283–284.

Schork, S.M., Thumm, M., and Wolf, D.H. (1995). Catabolite inactivation of fructose-1,6- bisphosphatase of Saccharomyces cerevisiae. Degradation occurs via the ubiquitin pathway. J Biol Chem 270, 26446–26450.

Schüle, T., Rose, M., Entian, K.D., Thumm, M., and Wolf, D.H. (2000). Ubc8p functions in catabolite degradation of fructose-1,6-bisphosphatase in yeast. The EMBO Journal 19, 2161–2167.

Schweitzer, A., Aufderheide, A., Rudack, T., Beck, F., Pfeifer, G., Plitzko, J.M., Sakata, E., Schulten, K., Forster, F., and Baumeister, W. (2016). Structure of the human 26S proteasome at a resolution of 3.9 A. Proc Natl Acad Sci U S A 113, 7816–7821.

Shakeel, S., Rajendra, E., Alcon, P., O’Reilly, F., Chorev, D.S., Maslen, S., Degliesposti, G., Russo, C.J., He, S., Hill, C.H., et al. (2019). Structure of the Fanconi anaemia monoubiquitin ligase complex. Nature 575, 234–237.

Skraban, C.M., Wells, C.F., Markose, P., Cho, M.T., Nesbitt, A.I., Au, P.Y.B., Begtrup, A., Bernat, J.A., Bird, L.M., Cao, K., et al. (2017). WDR26 Haploinsufficiency Causes a Recognizable Syndrome of Intellectual Disability, Seizures, Abnormal Gait, and Distinctive Facial Features. Am J Hum Genet 101, 139–148.

Soni, S., Bala, S., Gwynn, B., Sahr, K.E., Peters, L.L., and Hanspal, M. (2006). Absence of erythroblast macrophage protein (Emp) leads to failure of erythroblast nuclear extrusion. J Biol Chem 281, 20181–20189.

Tu, B.P., and McKnight, S.L. (2006). Metabolic cycles as an underlying basis of biological oscillations. Nat Rev Mol Cell Biol 7, 696–701.

Ulrich, A.K.C., Schulz, J.F., Kamprad, A., Schutze, T., and Wahl, M.C. (2016). Structural Basis for the Functional Coupling of the Alternative Splicing Factors Smu1 and RED. Structure 24, 762–773.

Watson, E.R., Brown, N.G., Peters, J.M., Stark, H., and Schulman, B.A. (2019). Posing the APC/C E3 Ubiquitin Ligase to Orchestrate Cell Division. Trends Cell Biol 29, 117–134.

Wehmer, M., Rudack, T., Beck, F., Aufderheide, A., Pfeifer, G., Plitzko, J.M., Forster, F., Schulten, K., Baumeister, W., and Sakata, E. (2017). Structural insights into the functional cycle of the ATPase module of the 26S proteasome. Proc Natl Acad Sci U S A 114, 1305–1310.

Wei, Q., Boulais, P.E., Zhang, D., Pinho, S., Tanaka, M., and Frenette, P.S. (2019). Maea expressed by macrophages, but not erythroblasts, maintains postnatal murine bone marrow erythroblastic islands. Blood 133, 1222–1232.

Weissman, J.S., Rye, H.S., Fenton, W.A., Beechem, J.M., and Horwich, A.L. (1996). Characterization of the active intermediate of a GroEL-GroES-mediated protein folding reaction. Cell 84, 481–490.

Welcker, M., Larimore, E.A., Swanger, J., Bengoechea-Alonso, M.T., Grim, J.E., Ericsson, J., Zheng, N., and Clurman, B.E. (2013). Fbw7 dimerization determines the specificity and robustness of substrate degradation. Genes Dev 27, 2531–2536.

Zaman, S., Lippman, S.I., Zhao, X., and Broach, J.R. (2008). How Saccharomyces responds to nutrients. Annu Rev Genet 42, 27–81.

Zavortink, M., Rutt, L.N., Dzitoyeva, S., Henriksen, J.C., Barrington, C., Bilodeau, D.Y., Wang, M., Chen, X.X.L., and Rissland, O.S. (2020). The E2 Marie Kondo and the CTLH E3 ligase clear deposited RNA binding proteins during the maternal-to-zygotic transition. eLife.

Zhen, R., Moo, C., Zhao, Z., Chen, M., Feng, H., Zheng, X., Zhang, L., Shi, J., and Chen, C. (2020). Wdr26 regulates nuclear condensation in developing erythroblasts. Blood 135.

Zhu, J., and Thompson, C.B. (2019). Metabolic regulation of cell growth and proliferation. Nat Rev Mol Cell Biol 20, 436–450.

Zhuang, M., Calabrese, M.F., Liu, J., Waddell, M.B., Nourse, A., Hammel, M., Miller, D.J., Walden, H., Duda, D.M., Seyedin, S.N., et al. (2009). Structures of SPOP-substrate complexes: insights into molecular architectures of BTB-Cul3 ubiquitin ligases. Mol Cell 36, 39–50.

