## supplementary_figures for "GID E3 ligase supramolecular chelate assembly configures multipronged ubiquitin targeting of an oligomeric metabolic enzyme"

### Supplemental Information

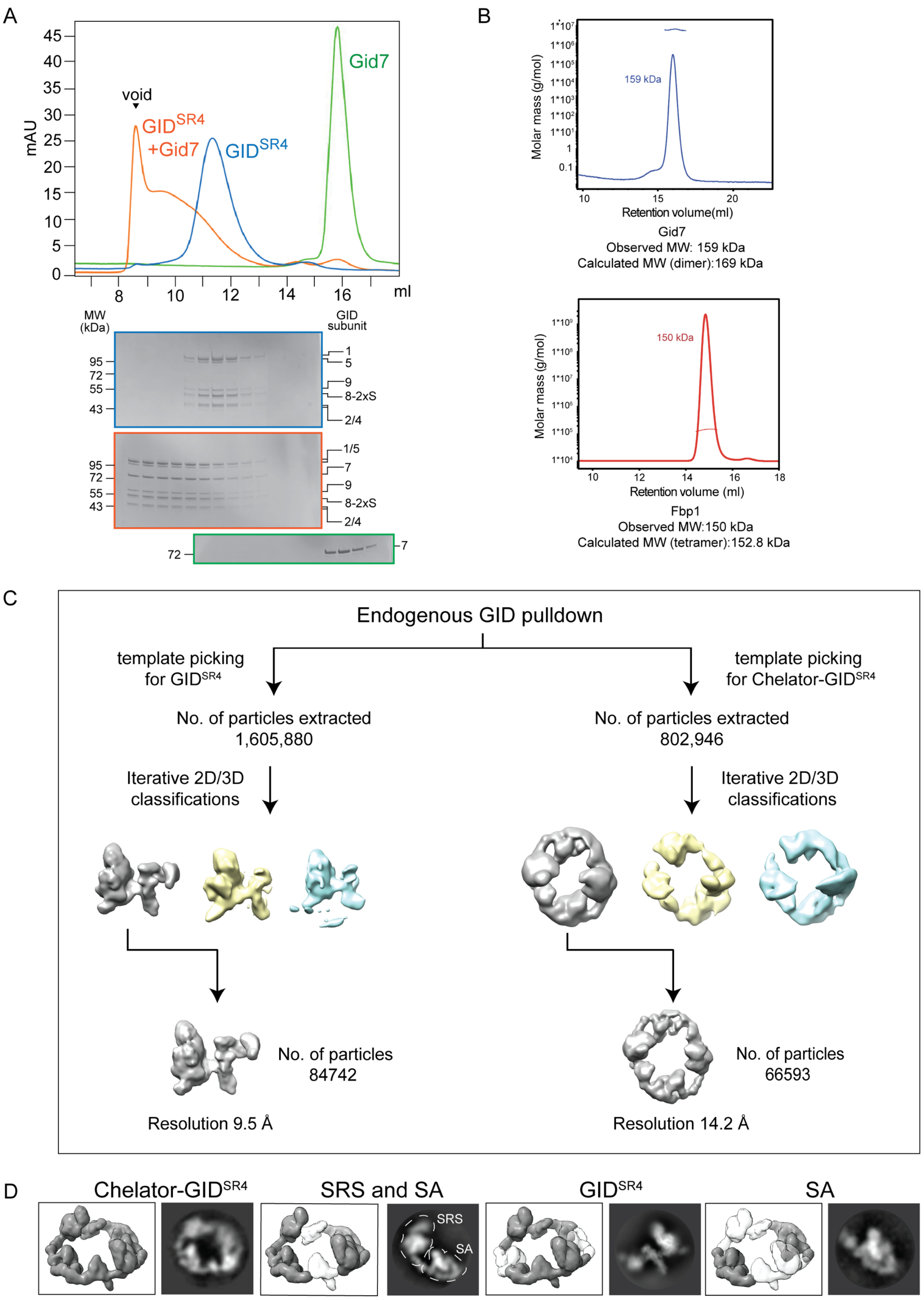

**Figure S1. Characterization of supramolecular assembly of Chelator-GID$^{SR4}$ (Related to Figure 1)**

A. Gel filtration analysis (top) of GID$^{SR4}$ (blue), GID$^{SR4}$ + Gid7 (orange) and Gid7 alone (green). Fractions after gel filtration were run on SDS-PAGE and visualized with Coomassie (bottom).
B. SEC-MALS testing the oligomeric state of Fbp1 (bottom) and Gid7 (top). By comparing the observed and calculated molecular weights, the tetrameric state of Fbp1 and dimeric state of Gid7 were revealed. The blue and red peaks show the absorption spectra at 280 nm for Gid7 and Fbp1, respectively, whereas the horizontal lines indicate their molar masses.
C. Flowchart of cryo EM processing of the endogenous yeast GID dataset.
D. Representative 2D classes originating from multiple cryo EM datasets showing stability of individual modules even in the absence of supramolecular assembly. The composition of each analysed sample is represented as gray density (to the left of each 2D class) in the map of Chelator-GID$^{SR4}$ (the white parts indicate the omitted modules).

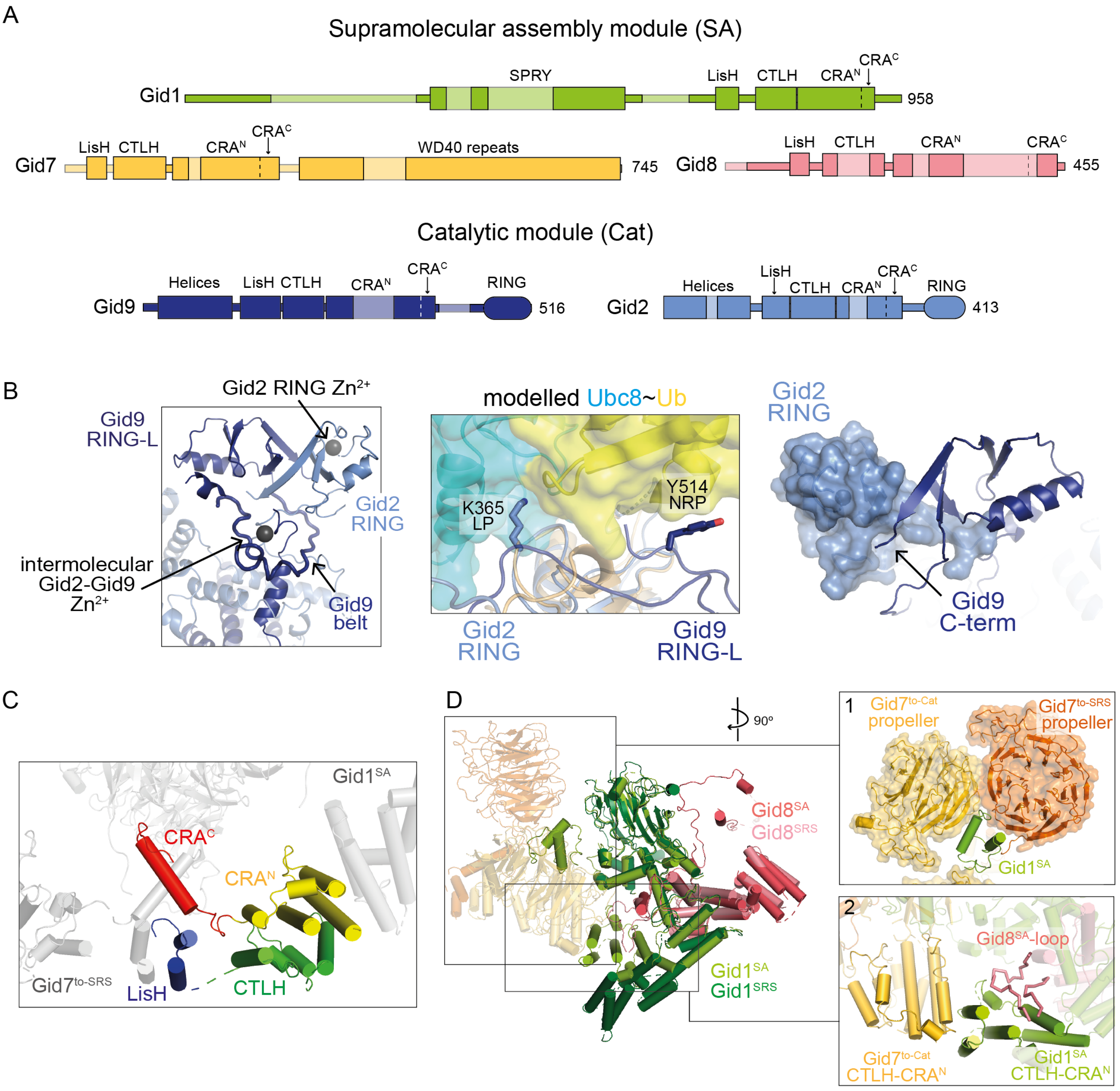


**Figure S2. Structural features of Chelator-GID$^{SR4}$ catalytic and supramolecular assembly modules (Related to Figure 2)**

A. Domain schematic of supramolecular assembly and catalytic modules. Darker regions represent parts of the subunits for which an atomic model was built.

B. Details of a novel heterodimeric Gid2-Gid9 RING-RING-like assembly (left) (zinc ions are represented as gray spheres). Gid2's RING domain binds only one zinc ion, in a manner reminiscent of SP-RINGs found in SUMO E3s. In contrast, Gid9 adopts an unconventional RING-like domain (RING-L) resembling U-box fold. The heterodimer is stabilized by an intermolecular zinc-binding domain as well as a Gid9 "belt". Model of E2~Ub activation by Gid2-Gid9 was generated by aligning the E2~Ub-bound RING structure (5H7S.PDB) with Gid2 RING (center). In the model, the previously characterized ‘linchpin’ residue (LP) of Gid2 and non-RING priming element (NRP) in Gid9 stabilize the closed conformation of Ubc8~Ub. The dimeric RING-RING-L assembly is stabilized by the packing of an extreme C-terminus of Gid9 against Gid2’s RING domain (shown as surface representation) (right).

C. Color-coded close-up view of LisH-CTLH-CRA dimerization domain in Gid7$^{to\text{-}Cat}$, where its most N- and C-terminal parts are colored blue and red, respectively. CRA is subdivided into N-terminal (CRA$^{N}$) and C-terminal (CRA$^{C}$) parts since they are components of 2 structurally and functionally distinct elements involved in intra- (LisH-CRA$^{C}$) and intermodular (CTLH-CRA$^{N}$) interactions.
D. Overlay of Gid1$^{SRS}$-Gid8$^{SRS}$ with Gid1$^{SA}$-Gid8$^{SA}$ in supramolecular assembly (SA) module (Gid7 dimer is kept transparent) together with close-ups highlighting: 1. a part of Gid1$^{SA}$ binding the asymmetric groove between the two Gid7 propellers and 2. SA-specific interactions between CTLH-CRA$^{N}$ of Gid1$^{SA}$, Gid8$^{SA}$ loop and CTLH-CRA$^{N}$ of Gid7$^{to\text{-}Cat}$.

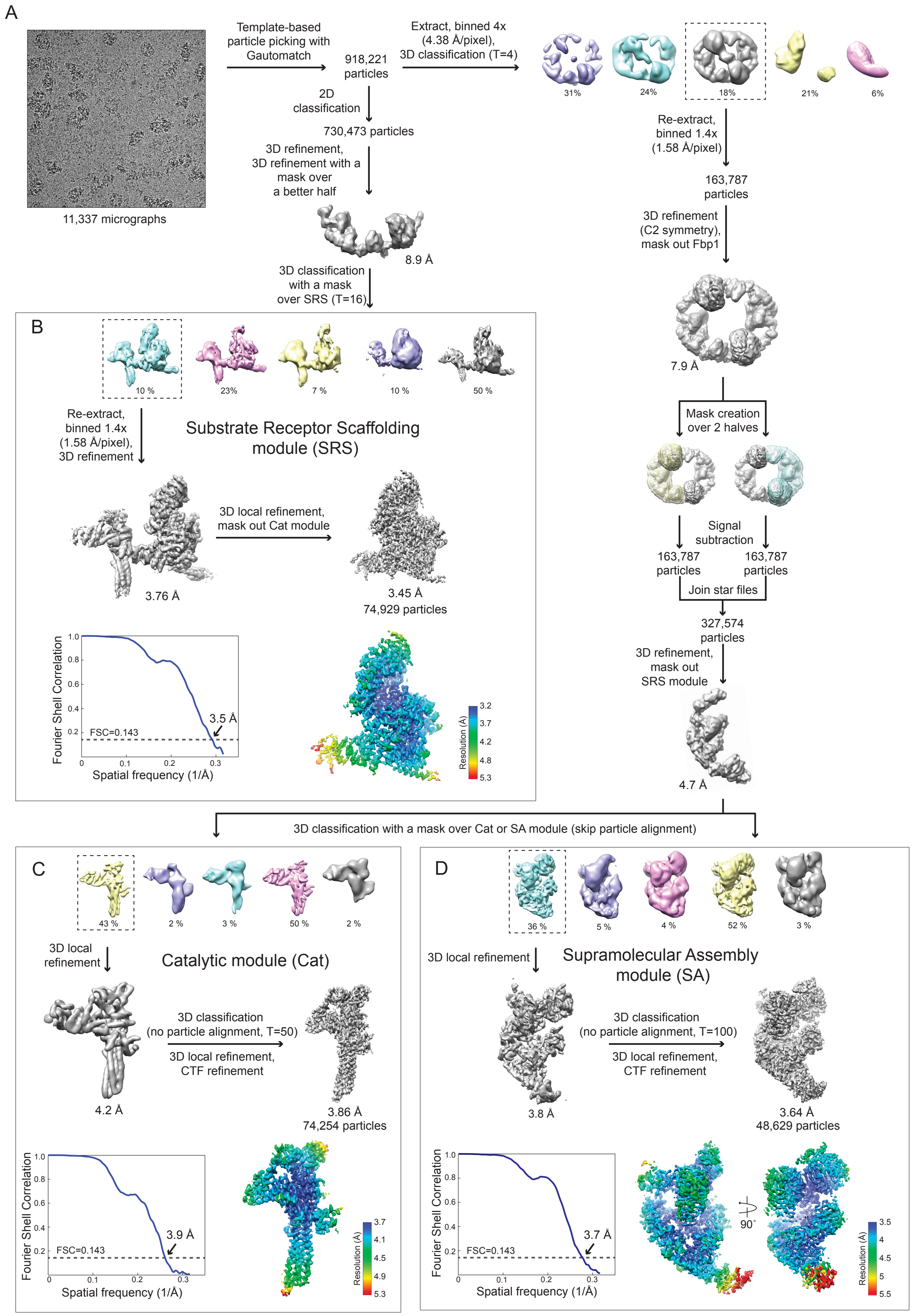
A
Template-based particle picking with Gautomatch
918,221 particles
Extract, binned 4x (4.38 Å/pixel), 3D classification (T=4)
31%
24%
18%
21%
6%
11,337 micrographs
2D classification
730,473 particles
3D refinement, 3D refinement with a mask over a better half
8.9 Å
3D classification with a mask over SRS (T=16)
Re-extract, binned 1.4x (1.58 Å/pixel)
163,787 particles
3D refinement (C2 symmetry), mask out Fbp1
7.9 Å
Mask creation over 2 halves
Signal subtraction
163,787 particles
163,787 particles
Join star files
327,574 particles
3D refinement, mask out SRS module
4.7 Å
B
10 %
23%
7 %
10 %
50 %
Re-extract, binned 1.4x (1.58 Å/pixel), 3D refinement
Substrate Receptor Scaffolding module (SRS)
3D local refinement, mask out Cat module
3.76 Å
3.45 Å
74,929 particles
Fourier Shell Correlation
FSC=0.143
3.5 Å
Spatial frequency (1/Å)
Resolution (Å)
3.2
3.7
4.2
4.8
5.3
3D classification with a mask over Cat or SA module (skip particle alignment)
C
43 %
2 %
3 %
50 %
2 %
3D local refinement
Catalytic module (Cat)
4.2 Å
3D classification (no particle alignment, T=50)
3D local refinement, CTF refinement
3.86 Å
74,254 particles
Fourier Shell Correlation
FSC=0.143
3.9 Å
Spatial frequency (1/Å)
Resolution (Å)
3.7
4.1
4.5
4.9
5.3
D
36 %
5 %
4 %
52 %
3 %
3D local refinement
Supramolecular Assembly module (SA)
3.8 Å
3D classification (no particle alignment, T=100)
3D local refinement, CTF refinement
3.64 Å
48,629 particles
Fourier Shell Correlation
FSC=0.143
3.7 Å
Spatial frequency (1/Å)
90˚
Resolution (Å)
3.5
4
4.5
5
5.5

**Figure S3. Simplified schematics for processing of high resolution cryo EM dataset of Chelator-GID$^{SR4}$ (Related to Figures 2, 3, 4)**

A. Flow chart of initial steps of processing yielding full and half maps of Chelator-GID$^{SR4}$
B. Focused 3D classification and local refinement of substrate receptor scaffolding module
C. Focused 3D classification and local refinement of catalytic module
D. Focused 3D classification and local refinement of supramolecular assembly module. For (B), (C) and (D), gold-standard Fourier shell correlation (FSC) curve (left bottom) and map of individual modules color-coded by their local resolution (right bottom) are shown. The dotted line in the FSC plot represents 0.143 cut-off criterion for nominal resolution.

A

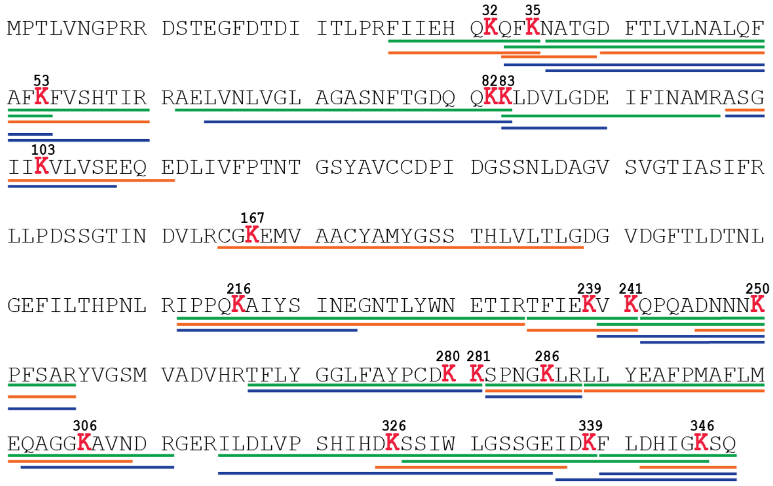


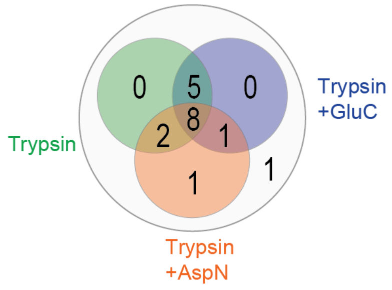


B

| Position | Sequence window | Localization probability | Intensity | Score | Score for localization |
|---|---|---|---|---|---|
| K32 | DTDIITLPRFIIEHQKQFKNATGDFTLVLNA | 1.00 | 1.43E+11 | 263.55 | 194.24 |
| K35 | IITLPRFIIEHQKQFKNATGDFTLVLNALQF | 1.00 | 5.33E+09 | 477.54 | 477.16 |
| K82 | LVGLAGASNFTGDQQKKLDVLGDEIFINAMR | 0.90 | 1.90E+08 | 239.58 | 239.58 |
| K83 | VGLAGASNFTGDQQKKLDVLGDEIFINAMRA | 1.00 | 2.66E+08 | 182.39 | 120.21 |
| K103 | GDEIFINAMRASGIIKVLVSEEQEDLIVFPT | 1.00 | 2.04E+07 | 188.26 | 188.26 |
| K216 | GEFILTHPNLRIPPQKAIYSINEGNTLYWNE | 1.00 | 2.83E+08 | 177.03 | 162.65 |
| K239 | GNTLYWNETIRTFIEKVKQPQADNNNKPFSA | 1.00 | 3.62E+07 | 195.57 | 195.57 |
| K280 | HRTFLYGGLFAYPCDKKSPNGKLRLLYEAFP | 1.00 | 9.97E+10 | 288.35 | 288.35 |
| K281 | RTFLYGGLFAYPCDKKSPNGKLRLLYEAFPM | 0.89 | 9.04E+10 | 161.65 | 97.797 |
| K326 | RGERILDLVPSHIHDKSSIWLGSSGEIDKFL | 1.00 | 7.26E+07 | 80.69 | 80.69 |

C

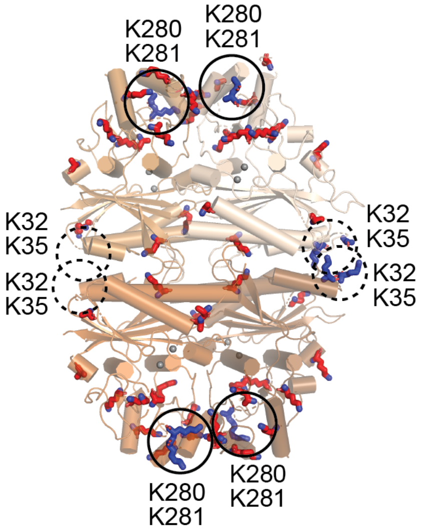


D

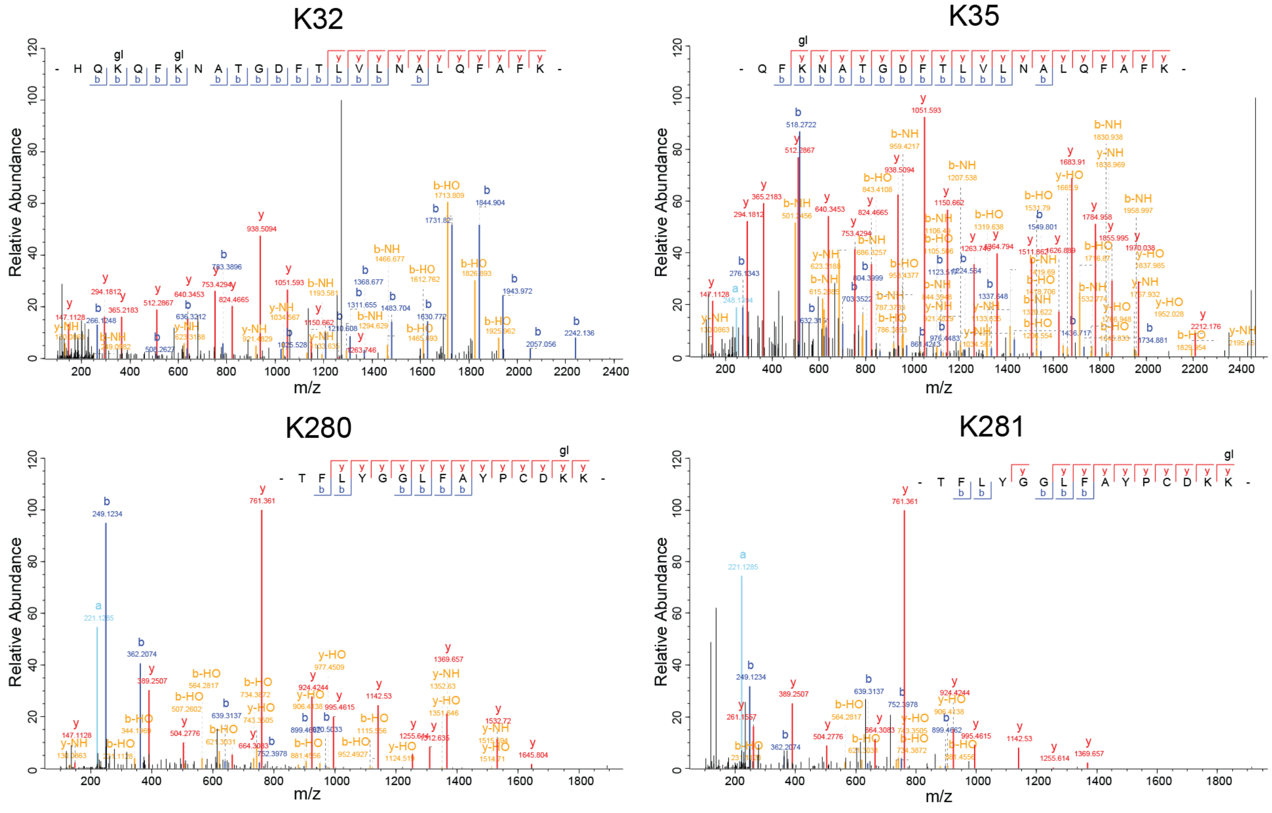

**Figure S4. Proteomic analysis disclosing preferred lysine sites on Fbp1 that are targeted for ubiquitylation by Chelator-GID$^{SR4}$ (Related to Figure 5)**

A. The positions in amino acid sequence (left, bold red letters) and numbers (right, shown as Venn diagram) of Fbp1 lysines that can be theoretically covered after proteolytic cleavages using trypsin (green), trypsin and AspN (red) or trypsin and GluC (blue).
B. Table showing experimentally identified Fbp1 peptides containing ubiquitylated lysine sites.
C. Crystal structure of Fbp1 tetramer with all the lysines shown as sticks. The most prominent ubiquitylation hits (which are highlighted in B), colored blue, are located in two clusters, K32/K35 (dashed circles) and K280/K281 (solid circles), in each Fbp1 protomer.
D. Tandem mass spectra ("best localization") for selected di-Gly remnant-modified peptides.

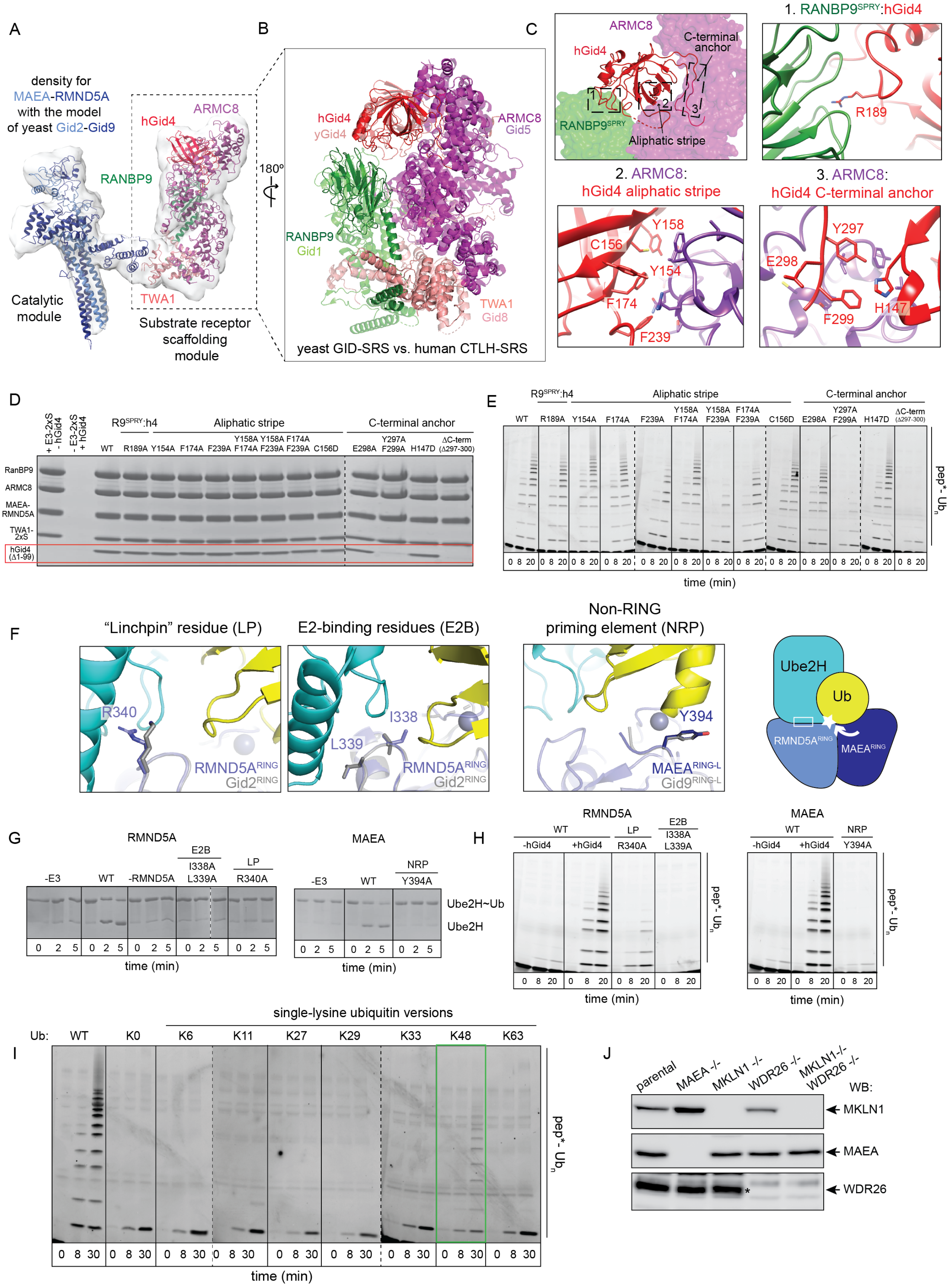
A
density for MAEA-RMND5A with the model of yeast Gid2-Gid9
hGid4
ARMC8
RANBP9
Catalytic module
TWA1
Substrate receptor scaffolding module
B
180º
hGid4
yGid4
ARMC8
Gid5
RANBP9
Gid1
TWA1
Gid8
yeast GID-SRS vs. human CTLH-SRS
C
ARMC8
C-terminal anchor
hGid4
RANBP9SPRY
Aliphatic stripe
1. RANBP9SPRY:hGid4
R189
2. ARMC8:
hGid4 aliphatic stripe
Y158
C156
Y154
F174
F239
3. ARMC8:
hGid4 C-terminal anchor
Y297
E298
H147
F299
D
+ E3-2xS - hGid4
- E3-2xS + hGid4
R9SPRY:h4
Aliphatic stripe
C-terminal anchor
WT R189A Y154A F174A F239A
Y158A F174A
Y158A F239A
F174A F239A
C156D E298A
Y297A F299A
H147D
ΔC-term (Δ297-300)
RanBP9
ARMC8
MAEA-RMND5A
TWA1-2xS
hGid4 (Δ1-99)
E
R9SPRY:h4
Aliphatic stripe
C-terminal anchor
WT R189A Y154A F174A F239A
Y158A F174A
Y158A F239A
F174A F239A
C156D E298A
Y297A F299A
H147D
ΔC-term (Δ297-300)
pep*- Ubn
0 8 20
time (min)
F
"Linchpin" residue (LP)
R340
RMND5ARING
Gid2RING
E2-binding residues (E2B)
I338
L339
RMND5ARING
Gid2RING
Non-RING priming element (NRP)
Y394
MAEARING-L
Gid9RING-L
Ube2H
Ub
RMND5ARING
MAEARING
G
RMND5A
-E3
WT
-RMND5A
E2B
I338A L339A
LP
R340A
0 2 5
time (min)
MAEA
-E3
WT
NRP
Y394A
Ube2H~Ub
Ube2H
H
RMND5A
WT
-hGid4
+hGid4
LP
R340A
E2B
I338A L339A
pep*- Ubn
0 8 20
time (min)
MAEA
WT
-hGid4
+hGid4
NRP
Y394A
I
single-lysine ubiquitin versions
Ub:
WT
K0
K6
K11
K27
K29
K33
K48
K63
pep* - Ubn
0 8 30
time (min)
J
parental
MAEA -/-
MKLN1 -/-
WDR26 -/-
MKLN1-/- WDR26 -/-
WB:
MKLN1
MAEA
WDR26
*

**Figure S5. Structural and mechanistic features of human CTLH (Related to Figure 6)**

A. Low resolution density of human CTLH$^{SR4}$ fit with atomic coordinates of CTLH SRS module and the yeast Cat module (Gid2-Gid9 heterodimer).
B. Overlay of the models of yeast GID (transparent) and human CTLH (opaque) substrate receptor scaffolding modules, showing structural homology.
C. Overview of major elements of hGid4 that enable its incorporation into CTLH$^{SR4}$. Close-ups of the key binding interfaces: 1. RANBP9$^{SPRY}$:hGid4 (denoted as R9$^{SPRY}$:h4 in assay panels), 2. ARMC8:hGID4 aliphatic stripe and 3. ARMC8:hGid4 C-terminal anchor. hGid4 residues that are mutated in (D) and (E) for biochemical assays are shown as sticks.
D. Assay testing the importance of hGid4 residues shown in (C) to its binding to the CTLH substrate receptor scaffolding module. The band corresponding to hGid4 in Coomassie-stained SDS-PAGE is highlighted with a red box. Binding of hGid4 to the complex was only impaired by mutations of its C-terminal anchor.
E. *In vitro* multi-turnover assays testing importance of hGid4 residues shown in (C) to ubiquitylation of a fluorescent model peptide substrate with a target lysine at position 27. Only the mutations of the key residues in the hydrophobic stripe of hGid4 $\beta$-barrel and its C-terminal anchor inhibited ubiquitylation.
F. Overlay of RMND5A (left) and MAEA (center) homology models with yeast Gid2-Gid9 RING-RING-L structure. Candidate "linchpin" (LP) and E2-binding (E2B) residues in RMND5A, and non-RING priming element (NRP) in MAEA are shown as sticks. These residues correspond to the LP K365 and E2B V363, L364 in yeast Gid2, and NRP Y514 in yeast Gid9 (shown as gray sticks). The significance of the LP, E2B and NRP in yeast Gid2-Gid9 for Fbp1 degradation *in vivo* have been shown previously in Qiao et al., 2020. Cartoon (right) summarizing the proposed catalytic mechanism of CTLH$^{SR4}$ based on the structural models and biochemical assays (shown in G and H). RMND5A is an active RING that directly binds its cognate E2 Ube2H (white square) and bears a "linchpin" residue (white star), whereas MAEA RING-L provides a non-RING priming element (white arrow).
G. Discharge assays showing effects of point mutations in RMND5A and MAEA as shown in (F) on intrinsic catalytic activity of CTLH$^{SR4}$. Discharge of Ube2H~Ub intermediate to free lysine in solution was detected with Coomassie-stained non-reducing SDS-PAGE.
H. *In vitro* multi-turnover assays showing effects of point mutations in RMND5A and MAEA as shown in (F) on ubiquitylation of a fluorescently labelled model peptide substrate (with a target lysine at position 27).
I. *In vitro* ubiquitylation assays with a panel of single lysine ubiquitin variants, as compared to WT and lysine-less (K0) ubiquitin, showing preference of CTLH$^{SR4}$ to form K48 polyubiquitin chains (highlighted in green box). The reactions were conducted with CTLH$^{SR4}$, E2 Ube2H and fluorescently labelled model peptide substrate (with target lysine at position 27).
J. Western blots confirming the MAEA, MKLN1, WDR26 and MKLN1/WDR26 knockouts in K562 cells. Asterisk indicates a WDR26 signal.

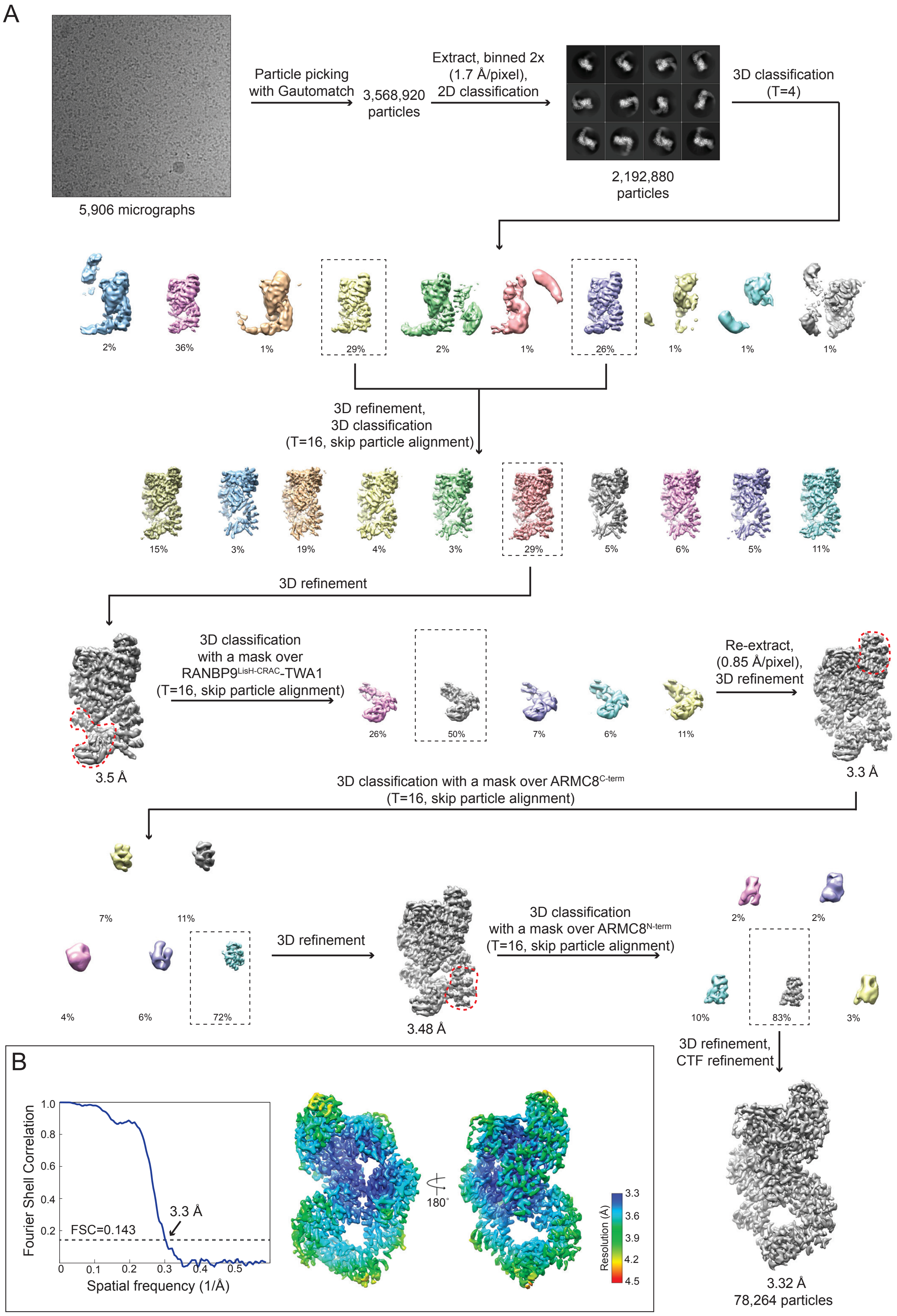
A
5,906 micrographs
Particle picking with Gautomatch
3,568,920 particles
Extract, binned 2x (1.7 Å/pixel), 2D classification
2,192,880 particles
3D classification (T=4)
2%
36%
1%
29%
2%
1%
26%
1%
1%
1%
3D refinement, 3D classification (T=16, skip particle alignment)
15%
3%
19%
4%
3%
29%
5%
6%
5%
11%
3D refinement
3.5 Å
3D classification with a mask over RANBP9LisH-CRAC-TWA1 (T=16, skip particle alignment)
26%
50%
7%
6%
11%
Re-extract, (0.85 Å/pixel), 3D refinement
3.3 Å
3D classification with a mask over ARMC8C-term (T=16, skip particle alignment)
7%
11%
4%
6%
72%
3D refinement
3.48 Å
3D classification with a mask over ARMC8N-term (T=16, skip particle alignment)
2%
2%
10%
83%
3%
3D refinement, CTF refinement
3.32 Å
78,264 particles
B
Fourier Shell Correlation
Spatial frequency (1/Å)
3.3 Å
FSC=0.143
180˚
Resolution (Å)
3.3
3.6
3.9
4.2
4.5

**Figure S6. Simplified schematics for processing of high resolution cryo EM dataset of human $CTLH^{SR4}$ (Related to Figure 6)**

A. Flow chart of cryo EM processing yielding a map of $CTLH^{SR4}$ substrate receptor scaffolding (SRS) module. 3D refinements were performed with a mask excluding the catalytic module due to its mobility relative to SRS. A series of focused 3D classification (with masks indicated as red dotted lines) was performed to resolve under-represented parts of the map.
B. Gold-standard Fourier shell correlation (FSC) curve (left) and the map of $CTLH^{SR4}$ SRS module color-coded by its local resolution (right). The dotted line in the FSC plot represents 0.143 cut-off criterion for nominal resolution.
